# Single cell resolved spatial immune repertoire unveils spatial heterogeneity of lymphoid aggregates in human immune disorders

**DOI:** 10.1101/2025.01.16.630222

**Authors:** Xiaojuan Zhan, Yi Liu, Yanying Guo, Wenwen Zhou, Yixin Yan, Hui Zeng, Xuan Dong, Xiaoyu Chen, Rong Ma, Zhong Liu, Fan Zhu, Xubin Zheng, Xinxing Li, Jinwen Yin, Francis Ka-ming Chan, Chuanyu Liu, Longqi Liu, Xun Xu, Yong Hou, Haoran Tao, Yuliang Dong, Tao Zeng, Young Li, Jingying Zhou, Zexian Zeng, Yu Feng

**Affiliations:** College of Life Sciences, University of Chinese Academy of Sciences, Beijing 100049, China; BGI Research, Shenzhen 518083, China; BGI Research, Hangzhou 310030, China; College of Life Sciences, Northwest University, Xi’an 710069, China; School of Computing and Information Technology, Great Bay University, Dongguan 523000, China; Guangdong Provincial Key Laboratory of Mathematical and Neural Dynamical Systems, Dongguan 52300, China; Zhejiang University Liangzhu Laboratory, 1369 West Wenyi Road, Hangzhou 311121, China; Department of Cardiology of The Second Affiliated Hospital, School of Medicine, Zhejiang University, Hangzhou 310009, China; State Key Laboratory of Transvascular Implantation Devices, Hangzhou 310009, China; Heart Regeneration and repair Key Laboratory of Zhejiang Province, Hangzhou 310009, China; Shenzhen Proof-of-Concept Center of Digital Cytopathology, BGI Research, Shenzhen 518083, China; Shanxi Medical University - BGI Collaborative Center for Future Medicine, Shanxi Medical University, Taiyuan 030001, China; Key Laboratory of Anti-Inflammatory and Immune Medicine, Ministry of Education, Anhui Medical University, Hefei 230032, China; Shenzhen Key Laboratory of Single-Cell Omics, BGI Research, Shenzhen 518083, China; School of Biomedical Sciences, The Chinese University of Hong Kong, Hong Kong SAR, 999077, China; BGI Hangzhou CycloneSEQ Technology Co., Ltd, Hangzhou 310030, China; Peking-Tsinghua Center for Life Sciences, Academy for Advanced Interdisciplinary Studies, Peking University, Beijing 100084, China; Center for Quantitative Biology, Academy for Advanced Interdisciplinary Studies, Peking University, Beijing 100084, China

## Abstract

Adaptive immunity, mediated by T and B cell responses, is essential for defending against infections and cancers while also being implicated in autoimmune diseases. Tracking T and B cell repertoires *in situ* at single-cell resolution is essential for understanding adaptive immune responses. To address the lack of tools for *in situ* single-cell T/BCR (XCR) sequencing, we developed Stereo-XCR-seq, an efficient strategy for retrieving and sequencing TCR and BCR from Stereo-seq cDNA libraries at subcellular resolution. Stereo-XCR-seq provides unbiased full-length XCR reads alongside spatial transcriptomics, enabling the identification of heterogeneous lymphoid aggregates with distinct clonal activities in cancers and inflammatory bowel disease (IBD). We identified plasma cell aggregates that differ from tertiary lymphoid structures (TLSs) in both transcriptomic profiles and clonal activities, with spatial positioning potentially mediating unique immune responses. Collectively, Stereo-XCR-seq enables *in situ* single-cell profiling of T and B cell clonal activities within tissue microenvironments, providing insights into lymphocyte adaption to environmental stimuli. This technology provides potential for advancing our understanding of tissue immunity and the development of therapeutic strategies for immune disorders.

## Introduction

The immune repertoire represents the diverse T-cell receptors (TCRs) and B-cell receptors (BCRs) that enable the immune system to recognize and respond to a wide range of antigens ^1^. This diversity arises during T and B cell maturation through variable, diversity, and joining (VDJ) recombination, producing unique and heritable TCR and BCR sequences ^2^. Upon antigen recognition, T and B cells undergo clonal expansion, a hallmark of immune activation that can be profiled through TCR and BCR sequencing. The composition and organization of the immune repertoire differ significantly across tissue contexts ^3–7^, highlighting the importance of studying immune repertoires within their spatial environment. In addition, spatial profiling of immune repertoires offers the potential to uncover how antigen-specific immune responses are orchestrated within tissues, providing critical insights into antigen recognition and immune system development ^8–10^. Therefore, the development of spatial immune repertoire technologies is a crucial step toward advancing our understanding of basic immunology and unlocking new clinical applications ^11^ .

Various toolkits have been developed to capture spatial immune repertoires *in situ* alongside transcriptomics, with notable examples including Slide-TCR-seq/Slide-tags ^12, 13^, Spatial VDJ ^14^, and SPTCR-seq ^15^. Slide-TCR-seq and Slide-tages ^12, 13^ leverages the Slide-seq platform^16^ for transcriptomics profiling and employ multiplexed PCR to enrich XCR transcripts. These methods enables the capture of both Coordinate ID (CID) and complementarity-determining region 3 (CDR3) sequences using short- or mid-read sequencing. However, its reliance on pre-designed PCR primer panels introduces potential biases stemming from prior knowledge. In contrast, SPTCR-seq ^15^ and Spatial VDJ ^14^ utilize the Visium platform for transcriptomics profiling and incorporate probes targeting the constant (C) region to enrich TCR transcripts through hybridization. While these approaches broaden accessibility to spatial immune repertoire profiling, they suffer from low efficiency, are time-consuming, and rely on hybridization-based enrichment methods that may limit sensitivity. Furthermore, these methods were designed for an earlier generation of the Visium platform with a spatial resolution of 55 µm, which lacks the subcellular precision now achievable with the latest advancements in spatial transcriptomics technologies.

Spatial transcriptomics has made significant strides, with commercial platforms now achieving subcellular resolution and high-throughput gene detection. Sequencing-based spatial transcriptomics technologies, such as Stereo-seq v1.3 ^17^ and Visium HD ^18^, have achieved resolutions of 0.5 μm and 2 μm, respectively. The rapid progress in these technologies underscores an urgent need to integrate them with spatial immune repertoire sequencing, offering a unique opportunity to gain deeper insights into the immune repertoire landscape and its functional organization within tissues ^8–10^. However, significant technical challenges remain in capturing TCR and BCR sequences *in situ* at single-cell resolution. First, the physical distances between VDJ regions and barcodes within transcripts exceed 1,000 base pairs, making them unsuitable for high-throughput short-read sequencing. Second, TCR and BCR transcripts are extremely rare, representing less than 0.01% of the cDNA library, which poses a challenge for low-throughput single-molecule long-read sequencing.

Upon the Stereo-seq platform ^17^, a technology we previously developed that combines DNA nanoball (DNB)-patterned arrays to capture poly-A mRNA at a resolution of 0.5 μm, we developed an efficient and rapid sequencing technology called single strain circle DNA PCR (sscirPCR) to retrieve TCR and BCR transcripts from the Stereo-seq cDNA library. This method, which we named Stereo-XCR-seq, enables unbiased immune repertoire profiling at single-cell resolution with paired-chain information in patient specimens from cancers and autoimmune diseases. Using Stereo-XCR-seq, we identified previously uncharacterized plasma cell aggregates that were distinct from tertiary lymphoid structures (TLSs) in terms of transcriptomic profiles and clonal activities. These aggregates exhibited spatial heterogeneity and were found in patients with lung and kidney cancers. Similarly, in patients with inflammatory bowel disease (IBD), mucosal plasma cells within these aggregates showed enhanced antigen-binding affinity and maturation activity, highlighting their potential role in disease pathology. The Stereo-XCR-seq platform represents a significant advance in the ability to perform *in situ* sub-cellular analysis of T and B cell clonal activities, offering new insights into immune dynamics and disease pathology.

## Results

### Stereo-XCR-seq profiles spatial T/BCR repertoires and transcriptomics at single-cell resolution

To achieve unbiased spatial immune repertoire profiling alongside subcellular spatial transcriptomics, we developed a spatial XCR sequencing strategy upon the Stereo-seq platform ^17^, a technology we previously developed (**Figure 1a**). We began by adding ligating elements (LE) to both ends of the Stereo-seq cDNA library through semi-nested PCR (**Figure 1a**, step 1) and circularizing it using XCR splint oligo-mediated ligation (**Figure 1a**, step 2). This process transformed the double-strained cDNA into end-to-end single-strained circular cDNA (sscirDNA), with barcodes ligated to the 5’ ends. Next, XCR transcripts were amplified via PCR using reverse primers targeting the constant (C) region (**Figure 1a**, step 3). This step remodeled the XCR transcripts into a new structure, with split C regions on both ends (C-ends) and the barcodes and VDJ regions in the middle.

**Figure 1.**
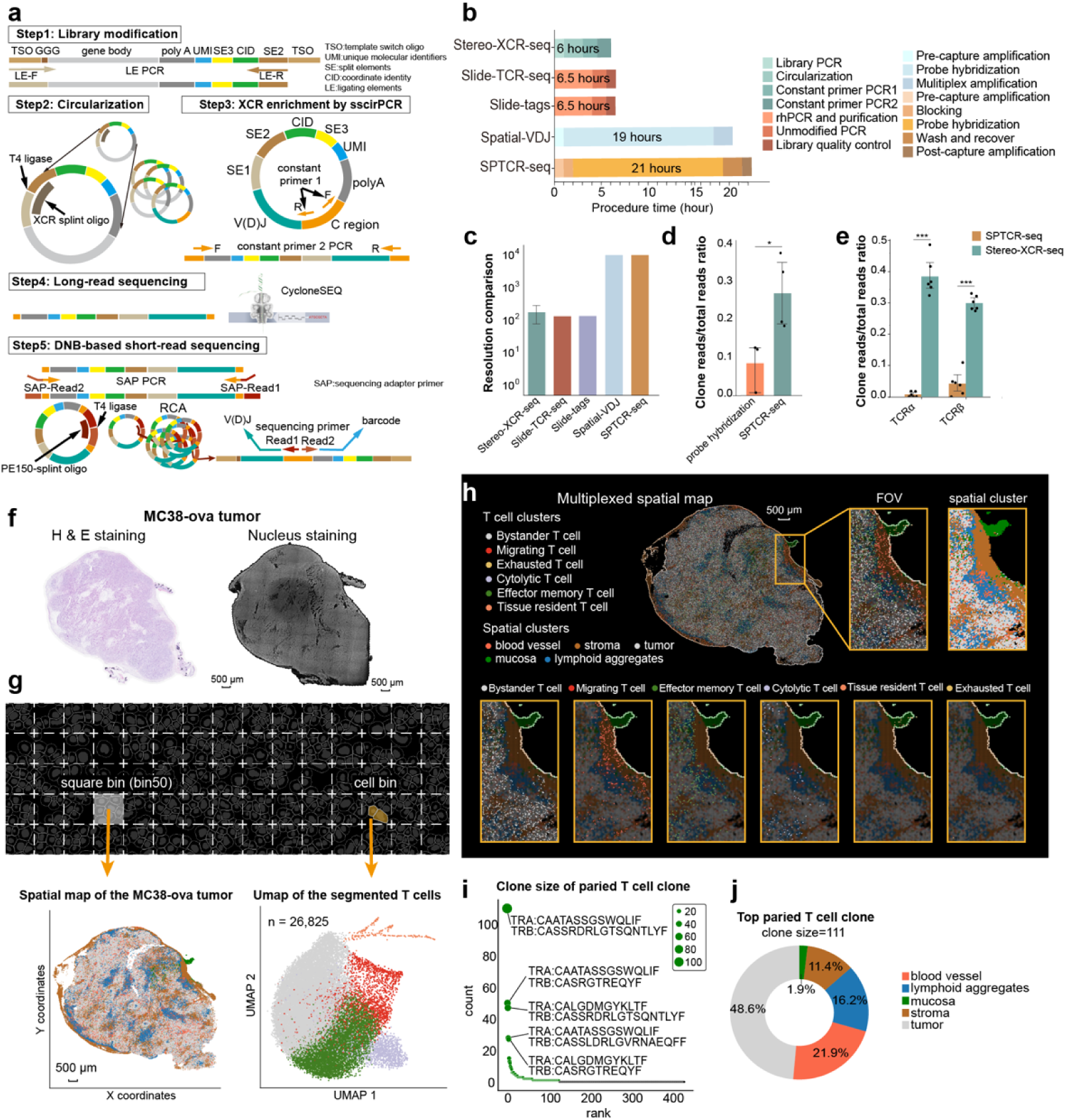
Stereo-XCR-seq profiles spatial T/BCR repertoires and transcriptomics at single-cell resolution. **a.** Schematic diagram of the workflow of Stereo-XCR-seq. The whole procedure can be divided into 5 steps, including library modification, circularization, sscirPCR, long-read sequencing, and short-read sequencing. Long-read sequencing and short-read sequencing are complementary, while either of them can generate coordinate barcoded clone reads. **b.** The bar chart shows the time used for each enrichment approach for spatial immune repertoire sequencing. The bars are colored by approaches and shaded by procedures. **c.** The bar chart shows the resolution of different technologies. Data are presented as median ± SD. **d.** The bar chart shows the enrichment efficiency of two unbiased enrichment approaches using stereo-seq cDNA libraries. Each dot represents an independent experimental replicate. N=3 for probe hybridization and N=4 for sscirPCR. Data are presented as median ± SD. **e.** The bar chart shows the data usage rate of stereo-XCR-seq and SPTCR-seq. The bars are colored by methodology. Data are presented as median ± SD. **f.** The H&E staining image of the adjacent tissue section and nucleus staining image of the tissue section on Stereo-seq chip of the MC38-ova tumor is shown (scale bar, 500 μm). **g.** The schematic diagram shows the generation of square bin (at a resolution of bin 50, 25 μm) and cell bin. The spatial map is made of filtered bin spots and colored by spatial clusters, showing the anatomical structure of the tumor (left, scale bar, 500 μm). The UMAP shows the cell clusters of single T cells segmented based on nucleus staining image (right). Each dot represents a T cell, colored by clusters and containing at least 1 TCRα or β clone. N cells= 26,825. **h.** The multiplexed spatial map was constructed with spatial map (underneath, bin50) colored by anatomical clusters and coordinate barcoded T cells (above, cell bin) colored by cell clusters. The fields of view highlight the distinguished distribution pattern across different T cell clusters. **i.** The scatter dot plot shows the clonal size of each paired clone. The dots are sized and ranked by clone size. N cells with paired TCRα and TCRβ chains = 974. **j.** The donut chart presents the distribution of T cells of the top paired clone. The percentage is labeled on the donut. N cells =111.

Long reads capture the full-length VDJ region and C region sequences, providing comprehensive coverage of the library construct, while short reads deliver higher throughput, cost-efficiency, and high-fidelity sequencing, enabling large-scale data generation and detailed analysis of somatic hypermutations. To balance accuracy and scalability, we combined both approaches to sequence the library construct. Half of the resulting library constructs in step 3 were processed with the CycloneSEQ platform^19^ to generate full-length long reads (**Figure 1a**, step4). The remaining constructs were truncated at both C-ends, and sequencing adapters (Read1 and Read2 primers) were added via nested PCR. These modified transcripts were then re-circularized for DNA nanoball-based paired-end 150 sequencing (PE150) to produce short reads (**Figure 1a**, step5). After sequencing, the raw long reads and short reads were processed separately based on their respective library structures (**Supplementary Fig.1a-b**) and subsequently integrated for downstream analyses. To mitigate the impact of sequencing errors and eliminate artificial clones, we retained only CDR3 clone types identified by short reads (**Supplementary Fig.1c**). For cell segmentation, a 500 nm (bin1) resolution mRNA image was manually registered to the nucleus staining image as previously described ^20^, and Cellpose V2 ^21^ was used to generate unique cell IDs and masks.

### Application and benchmarking of Stereo-XCR-seq for spatial immune repertoire profiling

We applied Stereo-XCR-seq to profile spatial transcriptomics and immune repertoires in tumor tissues from murine orthotopic colorectal tumor (CRC) models, patient samples with clear cell renal cell carcinoma (ccRCC) and non-small cell lung cancer (NSCLC), as well as mucosal biopsies from patients with inflammatory bowel disease (IBD). On average, we detected 44∼116 V genes, 20∼32 D genes, 22∼64 J genes of TCR, and 164∼171 V genes, 32 D genes, and 18 J genes of BCR in each specimen (**Supplementary Fig.2a**), demonstrating comprehensive coverage of VDJ genes. The VDJ regions were further assembled into 2,536∼16,910 CDR3 clones per sample (**Supplementary Fig.2b)**, with 25.09%∼66.58% of clones supported by both long reads and short reads (**Supplementary Fig.2c**). Notably, long reads provided broader coordinates coverage (62.35%∼94.34%) compared to short reads (8.67%∼48.47%, **Supplementary Fig.2d**), likely due to differences in whitelist mapping strategies.

We benchmarked Stereo-XCR-seq against multiple existing methods, including Slide-TCR-seq ^12^, Slide-tags ^13^, Spatial VDJ ^14^, and SPTCR-seq ^15^. Compared to these previously reported technologies for spatial immune repertoire profiling, Stereo-XCR-seq provides single-cell resolution, improved efficiency, and an unbiased enrichment strategy (**Figure 1b-c**, **Supplementary Fig.3a-c)**. Notably, Stereo-XCR-seq and Spatial VDJ are the only tools capable of capturing BCRs. Compared to Spatial VDJ ^14^, sscirPCR presented significantly higher enrichment efficiency over probe hybridization (fold change:3.164, **Figure 1d**) and led to a notable increase in data usage rate (**Figure 1e**). Compared to the existing tools, Stereo-XCR-seq identified a greater diversity of CDR3 clone types for each isotype and substantially higher lymphocyte cell number in non-lymphoid organs (**Supplementary Fig.3b-c)**. In addition, Stereo-XCR-seq is currently the only platform providing single-cell resolution with cellular morphology. By allocating clone reads to individual segmented cell bins, we observed pairing rates between 0.21% and 15.80% (**Supplementary Fig.3d**), with significantly higher rates for B/plasma cells (6.20% – 15.80%) compared to T cells (0.21% – 3.63%) (**Supplementary Fig.3e**). Together, these results highlight Stereo-XCR-seq’s superior performance and capture efficiency compared to existing tools ^12–15, 22^.

### High-resolution enables high-resolution mapping of clonal lymphoid cells

The Stereo-seq platform offers flexible spatial transcriptomics resolution, enabling simultaneous generation of square and cell bins on the same tissue section for studying clonal activities and anatomical structures ^23–25^. As a proof of concept, we applied Stereo-XCR-seq to an OVA-expressing MC38 tumor for data generation (**Figure 1f-g, Supplementary Fig.4a**). Using unsupervised clustering and transcriptomic markers, we identified multiple spatial structures, including tumor, stroma, lymphoid aggregates, blood vessels, and adjacent normal mucosa (analyzed at square bin50, ∼25μm x 25μm) (**Figure 1g, Supplementary Fig.4b-c**). A total of 26,825 T cells with at least one TCR clone read were identified, exhibiting high expression of T cell signatures (*Cd3d, Cd3e, Cd3g, Cd8a, Trac, Trbc1, Trbc2*) (**Figure 1h, Supplementary Fig. 4d**). Cell clustering revealed six T cell subsets (migrating, exhausted, cytolytic, effector memory, tissue-resident, and bystander T cells), representing 2,536 CDR3 clone types with distinct spatial distributions (**Supplementary Fig.4e**). These findings demonstrate Stereo-XCR-seq’s ability to map clonal lymphoid cells and their residing structures at high resolution.

Specifically, almost all tissue-resident T cells (*Cd69+, Itgae+*) were localized in the adjacent normal mucosa, over 60% of cytolytic T cells (*Gzma+, Gzme+*) were found within lymphoid aggregates, and more than 60% bystander T cells (no specific effector or memery markers) were found in the tumor region (**Figure 1h, Supplementary Fig.4f**). Among the 26,825 T cells identified, 974 were detected with paired TCRα and TCRβ chains, with clone sizes reaching up to 111 cells (**Figure 1i-j**). Notably, the top expanded paired clone (TCRα: CAATASSGSWQLIF, TCRβ: CASSRDRLGTSQNTLYF) ranked top for both α and β chains individually (**Supplementary Fig.4g**), indicating the accuracy of sequencing. These CDR3 sequences have previously been shown to specifically recognize the OVA-derived peptide SIINFEKL ^26^, further validating the accuracy of stereo-XCR-seq. Within the top expanded T cell clone, 87 cells (16%) were located in lymphoid aggregates, displaying strong stemness markers (*Tcf7, Il7r, Sell, Ccr7, Mki67*) and granzyme secretion activity (*Gzma, Gzmd, Gzmf, Gzmg, Gzmc, Gzmb*) (**Supplementary Fig.4h**). Together, our results suggest that the phenotype of the clonal T cells may be influenced by their spatial distribution within the tumor microenvironment.

### Stereo-XCR-seq profiles lymphoid aggregates in ccRCC tumor microenvironment

In renal cell carcinoma (RCC), both T and B lymphocytes contribute to anti-tumor immune response ^27, 28^, although their spatial clonal activities are rarely characterized. To address this gap, we applied Stereo-XCR-seq to analyze a tumor tissue sample from a patient with clear cell RCC (ccRCC) (**Figure 2a**). In the spatial transcriptome, we detected a median of 1,862 genes and 4,289 UMIs per analysis spot (bin50) (**Figure 2b**, **Supplementary Fig.5a**). For the spatial immune repertoire, we identified 1,831 TCR CDR3 clones and 7,317 BCR CDR3 clones (**Figure 1b**). Among the 35,110 genes detected in the spatial transcriptome, *immunoglobulin lambda* (*IGL*) and *immunoglobulin kappa* (*IGK*)-related genes exhibited the highest spatial autocorrelation, as indicated by their top-ranked Moran’s indices (**Supplementary Fig.5b**). This finding was further corroborated by the clustered expression of *IGL* transcripts and IgL clone reads in the spatial map (**Figure 2c-d**). A strong correlation (Pearson’s r = 0.76, p < 0.0001, **Figure 2e**) between the transcriptomic read counts and clone read counts provided robust evidence of the CIDs’ consistency after sscirPCR enrichment.

**Figure 2.**
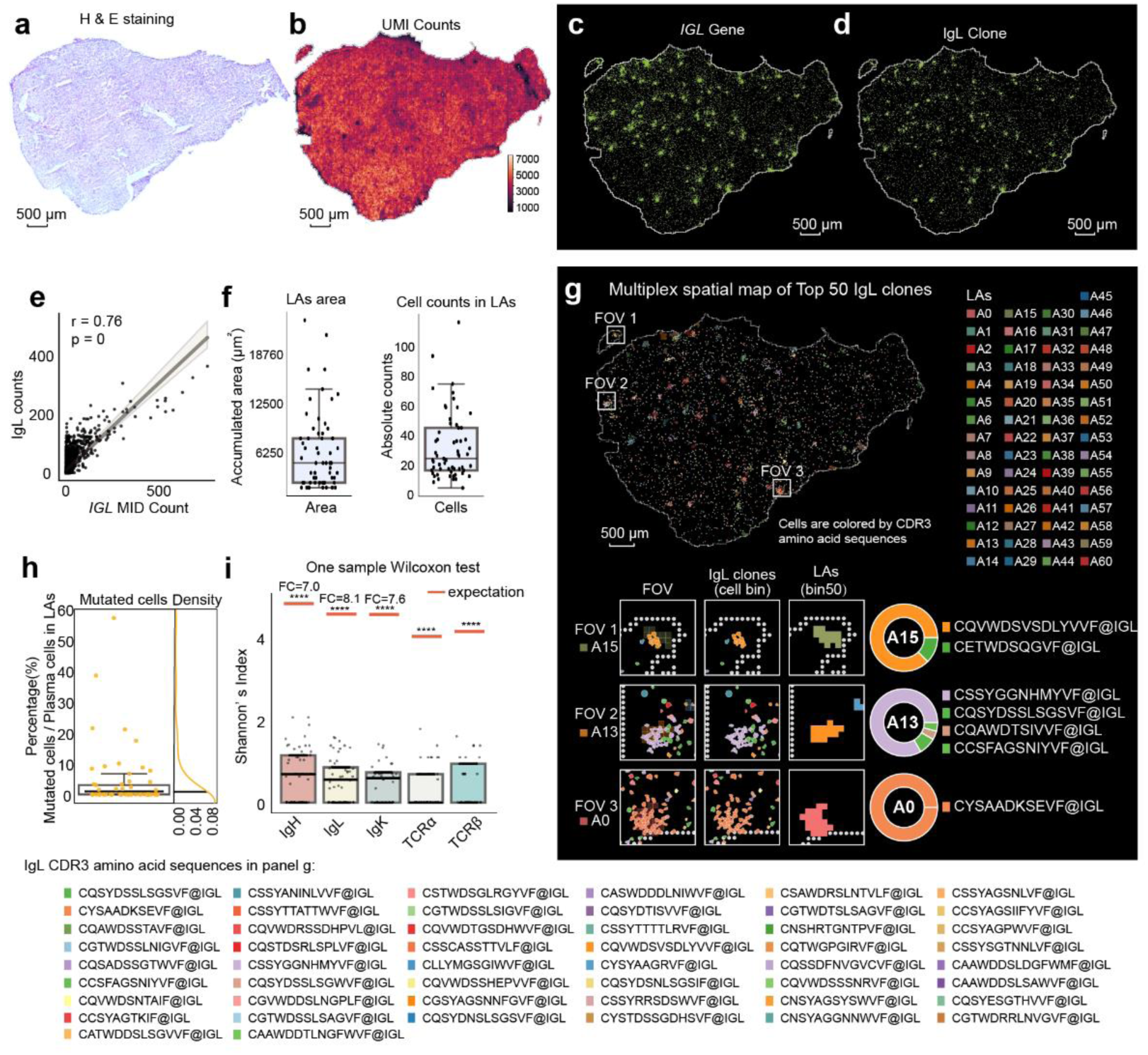
Stereo-XCR-seq profiles lymphoid aggregates in ccRCC tumor microenvironment. **a.** H&E staining of the ccRCC tumor (scale bar, 500 μm). **b.** The spatial plot shows the UMI counts of the spatial transcriptome (scale bar, 500 μm). Each dot represents a square bin (at a resolution of bin50). The dots are colored by UMI abundance. **c-d.** The spatial plot shows the spatial distribution of *IGL* transcripts (**c**) and IgL clones (**d**) at a resolution of 500 nm (bin1). Dots with at least one *IGL* transcript (**c**) or one IgL clone read (**d**) are colored in green (scale bar, 500 μm).**e.** The scatter plot and fitting line show the Pearson’s correlation between the abundances of the *IGL* transcripts (**c**) and the IgL clone (**d**) reads in the ccRCC tumor. The analyses were performed using the UMI counts of IGL transcripts and IgL clone read counts in each square bin at a resolution of bin50. The P-value and r-value are labeled above. **f.** The box-whisker plots show the area of the lymphoid aggregates (left) and cell counts in each lymphoid aggregate (right). Data are presented as median ± IQR (25%∼75%) with extreme values indicated by lower and upper bars. The outliers are indicated as hollow dots. **g.** The multiplexed spatial map shows the distribution of the top 50 IgL clones (at a resolution of CB) and lymphoid aggregates (at a resolution of bin50). The spatial plot of the lymphoid aggregates, colored by lymphoid aggregate labeling, is embedded underneath, overlain by the B/plasma cells colored by CDR3 amino acid sequences. The tissue outlines are highlighted by white dots. The FOVs show the representative lymphoid aggregates and the IgL clonal cells inside each lymphoid aggregate. The proportion and the clone size of each clone are shown in the corresponding donut plots. **h.** The box-whisker plot shows the percentage of mutated plasma cells (versus the total plasma cells) in each lymphoid aggregate. Data are presented as median ± IQR (25%∼75%) with extreme values indicated by lower and upper bars. The black line indicates the proportion of mutated plasma cell (versus the total plasma cells) outside the lymphoid aggregates as the expectation. Fold change is calculated using the median value versus the expectation. **i.** The box-whisker plot shows the Shannon’s Indices to indicate the clonal diversities of IgH, IgL, IgK, TCRα, and TCRβ clones in each lymphoid aggregate. The red lines indicate the Shannon’s Indices of each isoform outside the lymphoid aggregates as the expectation. Fold changes are calculated using median values versus expectations. The comparisons between the tumor and the lymphoid aggregates are performed using a one-sample Wilcoxon test.

To investigate the clonal activities of B/plasma cells in this tumor, we stratified IgL clones into four categories based on clone size: hyperexpanded (clone sizes≥20), medium (6≤clone sizes<20), small (2≤clone size<6), and not expanded (clone size=1). As expected, hyperexpanded clones displayed a highly clustered distribution with significant abundance, in contrast to the more dispersed pattern of smaller clones (**Supplementary Fig.5c**), as shown in the spatial plot and cell-to-cell distances for each clone (**Supplementary Fig.5d-e**). These observations suggest that the clonal activity of B/plasma cells may play a key role in driving the formation of lymphoid aggregates within the ccRCC tumor microenvironment.

To investigate the functional roles of these lymphoid aggregates, we denoised the BCR expression using the KDTree^29^ algorithm and identified 61 geographically discrete lymphoid aggregates through density-based spatial clustering (**Supplementary Fig.6a-d**). The median size of these lymphoid aggregates was 5,000 μm^2^ (interquartile range 25%∼75%: 2,500μm^2^∼8,125μm^2^) (**Figure 2f, left**), with a median of 21 cells per aggregate (interquartile range 25%∼75%: 13 ∼ 42 cells) (**Figure 2f, right**). These sizes were smaller than the minimal resolution (10,000 μm^2^) reported in recent studies ^14, 15^, but comparable to previously described glomerular IgA deposits in nephritis patients ^30^. To assess clonality within each aggregate, we quantified the cell numbers of each IgL clone and revealed that the majority of the lymphoid aggregates (45 out of 61) were dominated by a single clone, with clone sizes exceeding 50% of the total cell population (**Figure 2g, Supplementary Fig.6e**). Notably, 18 lymphoid aggregates contained only a single clone type, suggesting a high degree of clonal exclusivity in these regions.

We next examined whether lymphocytes primed in lymphoid aggregates were associated with antigen stimulation by examining somatic hypermutation, a hallmark of antigen-driven clonal activity for affinity maturation ^31, 32^. Only a small fraction of B/plasma cells (0.95%) outside lymphoid aggregates in the tumor exhibited mutated germline clones (**Figure 2h**, black line). In contrast, 21 lymphoid aggregates showed elevated proportions of cells with mutated IgL clones (median=6.67%, min=1.38%, max=57.14%) (**Figure 2h**), suggesting enhanced somatic hypermutation within these structures. Shannon indices revealed lower diversity of BCR chains within aggregates compared to tumor region (**Figure 2i**), indicating increased clonality. Intriguingly, although TCR indices suggested locoregional clonal expansions (**Figure 2i**), T cells were distributed more diffusely compared to aggregated B/plasma cells (**Supplementary Fig.7a-f**), with lower Moran ’ s indices and greater cell-to-cell distances (**Supplementary Fig.7g-h**). This scattering pattern may result from T cell emigration into tumor regions driven by *CXCL9/10* chemoattraction (**Supplementary Fig.7i-l**), the T cell trafficking role of which was previously reviewed in RCC ^33^. These findings suggest that micro-lymphoid aggregates in the ccRCC tumor drive antigen-driven immune responses by priming T and B/plasma cells, underscoring their critical role in shaping the immune response.

### Stereo-XCR-seq reveals spatial dynamics of B cell clonal activities in NSCLC lymphoid aggregates

B cells undergo somatic hypermutation to enhance antigen-binding affinity and class-switching recombination (CSR) to produce antibodies of distinct classes (IgA/G/E/D/M), a process typically occurring in germinal centers alongside T cell priming ^34–36^. To explore how these processes operate in the tumor microenvironment, we applied Stereo-XCR-seq to an NSCLC tumor with multiple lymphoid aggregates (**Figure 3a, Supplementary Fig.8-9**). Using unsupervised clustering ^37, 38^, we identified ten spatial structures (**Figure 3a, Supplementary Fig.8a**), including tertiary lymphoid structures (TLSs) and plasma cell aggregates (**Figure 3a, Supplementary Fig.8b-f**). Additionally, alveolar cells were observed encircling cancer cells, forming a clear peritumoral border (**Figure 3b**). Within the TLS clusters, five geographically discrete TLSs were identified (**Figure 3c**), including four peritumoral TLSs along the tumor border and one intratumoral TLS located within the tumor center. Comparisons of CDR3 clone types across these TLSs revealed that more than 60% of clones were unique to individual TLSs (**Figure 3d**), while fewer than 5% of CDR3 clones were shared across all five TLSs (**Supplementary Fig.9a-e**). These findings highlight substantial spatial heterogeneity among TLSs within the tumor microenvironment.

**Figure 3.**
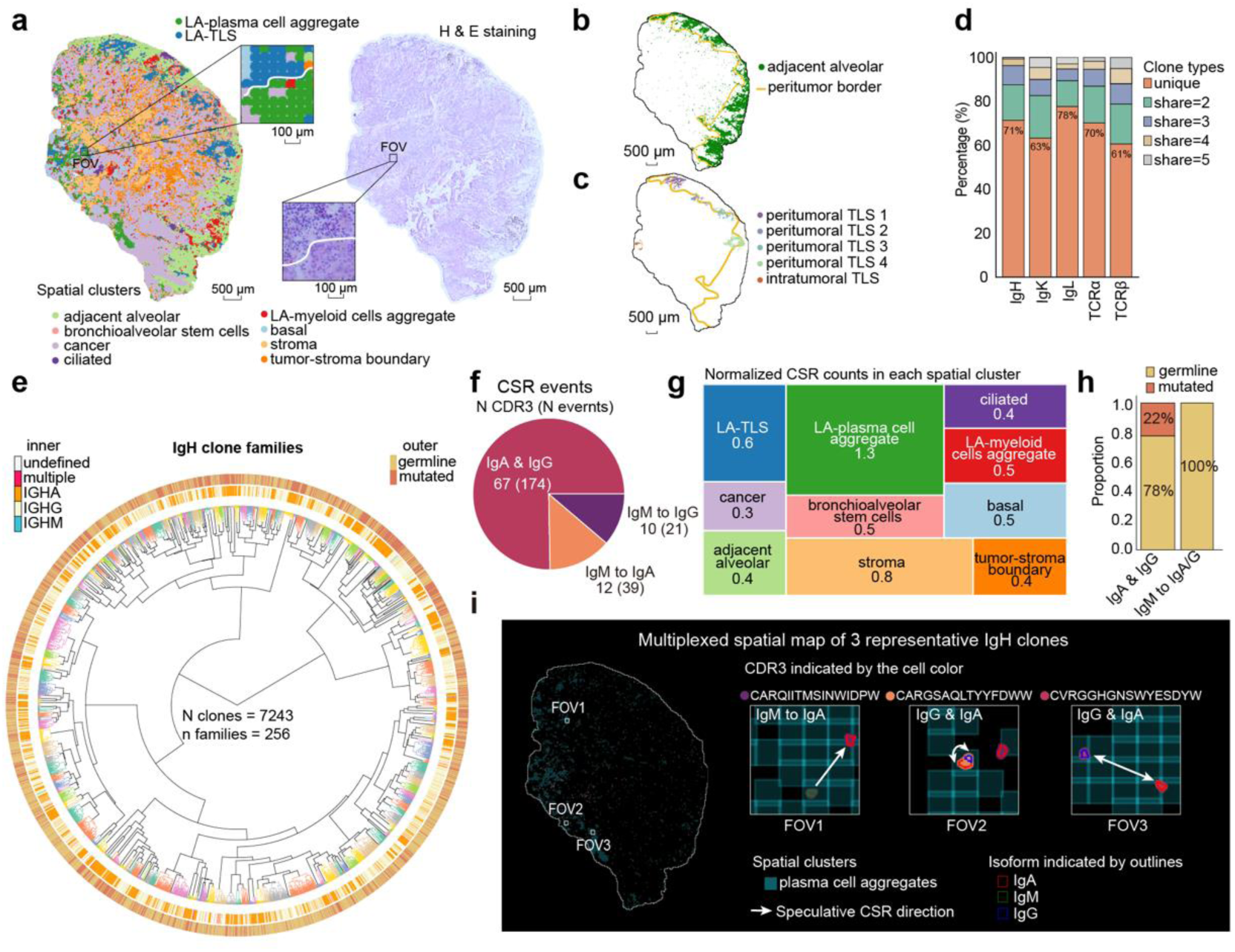
Stereo-XCR-seq reveals spatial dynamics of B cell clonal activities in NSCLC lymphoid aggregates. **a.** The spatial plot (left) and H&E staining (right) show the anatomic structure of the NSCLC tumor tissue (scale bar, 500 μm). Each dot in the spatial plot represents a square bin (at a resolution of bin50), colored by spatial clusters as indicated below. Two representative FOVs show the different anatomic structure of LA-plasma cell aggregate and LA-TLS (scale bar, 100 μm). **b**. The spatial plot shows peritumoral border (yellow fitting curve) and the adjacent normal alveolar tissue (scale bar, 500 μm). Each dot in the spatial plot represents a square bin (at a resolution of bin50), colored by spatial clusters as indicated below. **c.** The spatial plot shows peritumoral border (yellow fitting curve) and 5 geographically discrete TLSs (scale bar, 500 μm). Each dot in the spatial plot represents a square bin (at a resolution of bin50), colored by TLS labeling as indicated below. **d.** The stacked bar plot shows the sharing of clone types across 5 TLSs. **e.** The radial tree plot shows the lineage inference by hierarchical clustering of the IgH clones of the NSCLC tumor. Each terminal branch represents a clone type, colored by family. The inner circle is composed of lines indicating if this clone is a germline or mutated clone. The outer circle is composed of lines indicating the IgH classes. N clones = 7243, n families = 256. **f.** The pie chart shows the type of CSR events. A&G stands for the event that both IgA and IgG classes of an identical CDR3 clone could be detected in the same spatial cluster. IgM to IgA and IgM to IgG suggested the events both IgM and IgA or IgG of an identical CDR3 clone could be detected in the same spatial cluster. **g.** The tangram plot shows the frequencies of CSR events in each spatial cluster. The area of each rectangle is sized by the frequencies (normalized by the area of each spatial cluster). **h.** The stacked bar plot shows the proportion of mutated clones and germline clones in different CSR events. The stacked bars are colored by mutation status. **i.** The multiplexed spatial map shows CSR events of 3 representative IgH clones. The spatial plot of the plasma cell aggregates is embedded underneath, overlain by the B/plasma cells. The cells are filled by 3 different colors indicating 3 clones, with the outlines colored by classes. Three representative FOVs are selected to show the 3 different CSR events: 1. CSR of a germline clone (CDR3: CARQIITMSINWIDPW) from IgM to IgA. 2. CSR of a germline clone (CDR3: CVRGGHGNSWYESDYW) from IgG to IgA or from IgA to IgG. 3. CSR of a mutated clone (CDR3: CARGSAQLTYYFDWW) from IgG to IgA or from IgA to IgG.

The local activation of tumor-infiltrating lymphocytes in TLS promotes antitumor immunity ^39^, as evidenced by increased immune infiltration in TLS-positive tumors ^5,40^. However, the correlation between lymphoid aggregates and patient prognosis varies, with intratumoral TLSs associated with better outcomes than peritumoral TLSs ^3, 5, 41,42^. Given the role of TLSs in driving antigen-binding affinity maturation of B/plasma cells ^28^, we hypothesized that intratumoral TLSs exhibit enhanced maturation activity. To test this, we clustered 7,243 IgH clones into 256 families (**Figure 3e**), and categorized them as germline or somatically mutated based on nucleotide mutations. Notably, intratumoral TLSs showed significantly higher proportions of both germline (3.45% vs. 0.89∼1.92%) and mutated clones (1.38% vs. 0.22∼0.65%) compared to peritumoral (**Supplementary Fig.9f-g**). Analysis of CDR3 amino acid sequences using Levenshtein distances revealed strong autocorrelation among 48 IgH clones in the intratumoral TLS, with 47 mutated clones derived from 1 germline clone (**Supplementary Fig.9h**). Together, these findings highlight robust antigen-binding affinity maturation within intratumoral TLSs.

To delineate the sequential order of CSR and somatic hypermutation in B/plasma cells, we defined CSR events as the co-existence of at least 2 isotypes (IgM, IgG, or IgA) in the same spatial cluster. A total of 234 CSR events involving 89 IgH clones were identified, including 67 clones with both IgG and IgA isoforms, 10 clones transitioning from IgM to IgG, and 12 from IgM to IgA (**Figure 3f**). Plasma cell aggregates exhibited higher CSR frequencies compared to TLSs and other spatial clusters (**Figure 3g**), suggesting that plasma cell aggregates may serve as the primary niche for determining B/plasma cell effector functions, aligning with a recent study ^43^. Notably, 78% of the IgG/IgA clones were germline, and all IgM-to-IgG/A transitions occurred exclusively in germline clones (**Figure 3h**), implying that CSR can precede affinity maturation. Spatial mapping further validated this finding by identifying a germline clone transitioning from IgM to IgA/G (CDR3: CARQIITMSINWIDPW), a germline clone switching between IgA and IgG (CDR3: CVRGGHGNSWYESDYW), and a mutated clone transitioning between IgA and IgG (CDR3: CARGSAQLTYYFDWW) (**Figure 3i**). Collectively, these results demonstrated the capacity of Stereo-XCR-seq to delineate the clonal activities of lymphocytes in the lymphoid aggregates.

### Stereo-XCR-seq reveals the origins and clonal dynamics of lymphoid aggregates in IBD

The formation of lymphoid aggregates is frequently observed in organ-specific autoimmune diseases, including IBD ^44, 45^, though their correlation with disease activity remains unclear. This ambiguity may stem from inconsistent definitions of lymphoid aggregates and the elusive origins of disease-associated clones ^46^. To address this, we collected two mucosal biopsies from a patient with Crohn’s disease, one from mildly inflamed normal tissue and one from inflamed tissue (**Figure 4a-c**). Using spatial transcriptomic data and unsupervised clustering, we identified multiple spatial clusters, including lamina propria (*SELENOP*, *APOC3,* and *IL32*), ciliated epithelia (*DEFA5*, *DEFA6,* and *REG3A),* and three lymphoid aggregates (**Figure 4a, Supplementary Fig.10a**). A TLS marked by *CXCL13*, *CR2,* and *CCL19* was exclusively found in the submucosal region of inflamed tissue (**Supplementary Fig.10b**), consistent with its known prevalence in Crohn’s disease ^47^. Additionally, plasma cell aggregates – both central and peripheral, were larger and more prominent in the inflamed tissue compared to the normal counterpart (**Supplementary Fig.10c-d**). These findings suggest that lymphoid aggregates, particularly TLSs, and plasma cell aggregates, expand and localize differently in response to inflammation in Crohn’s disease.

**Figure 4.**
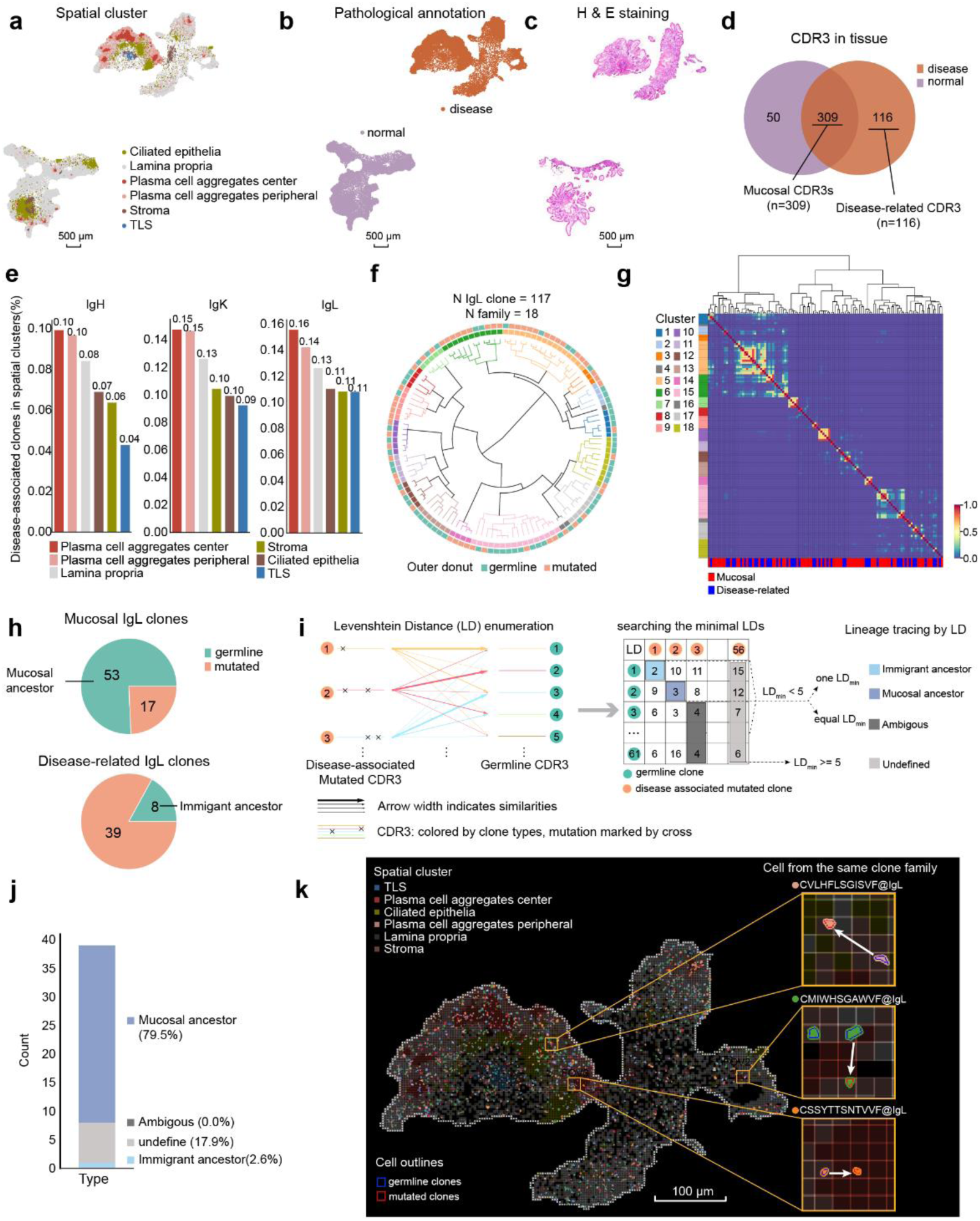
Stereo-XCR-seq reveals the origins and clonal dynamics of lymphoid aggregates in IBD. **a-c.** The spatial plot **(a-b)** and H&E staining (**c**) of the 2 IBD biopsies show the spatial clusters (**a**), pathological annotation (**b**), and anatomic structures (**c**) (scale bar, 500 μm). Each dot in the spatial plot represents a square bin (at a resolution of bin50), colored by spatial clusters or pathological annotation as indicated. **d.** The Venn diagram shows the shared and unique clones of the normal tissue and inflamed disease tissue. Only TCR and BCR chains allocated to cell segmentations were counted. **e.** The bar plot shows the proportion of disease-associated clones in each spatial cluster. The proportions are counted using IgH, IgK and IgL respectively. **f.** The radial tree plot shows the lineage inference by hierarchical clustering of the IgL clones of the inflamed tissue. Each terminal branch represents a clone type, colored by family as shown in the inner circle. The outer circle is composed of lines indicating if this clone is a germline or mutated clone. N clones = 117, n families = 18. **g.** The heatmap shows the pairwise CDR3 similarities. Each matrix is colored by the inferred similarity. The colored lines on the Y-axis are colored by the family. The colored lines on the below indicate if this clone is disease-related or shared by 2 biopsies. The tree plots on the above are identical to indicate the lineage inference of the IgL clones. **h.** The pie charts show the proportion and absolute number of mutated and germline IgL clones within shared and disease-related partitions. **i.** The schematic diagram shows the lineage tracing methodology. **j.** The stacked bar plot shows the origins of the disease-related mutated IgL clones. **k.** The multiplexed spatial map shows the distribution of disease-related IgL clones. The spatial plot showing the spatial clusters is embedded underneath, overlain by the B/plasma cells colored by CDR3 amino acid sequences. The outlines of cells indicate if this cell is of mutated (red) or germline (blue) clones. The FOVs show the juxtapositions of mutated cells and germline cells from the same clone family, with the CDR3 amino acid sequences labelled accordingly. Only cells of the indicated clone family are presented in the FOVs (scale bar, 100 μm).

To investigate the origins of disease-associated lymphocytes, we allocated TCR/BCR reads to cell barcodes and compared clonal patterns between normal and inflamed tissues. In total, we identified 425 T cells and 11,474 B/plasma cells (**Supplementary Fig.11a**), annotating 309 shared clones as mucosal and 116 clones as disease-related due to their absence in normal tissue (**Figure 4d**). A strong correlation in mucosal clone clonalities between the two tissues suggested similar antigen exposures (Pearson’s r = 0.86, p < 0.0001) (**Supplementary Fig.11b**), likely associated with the microbiome ^48^.

Interestingly, while all TLS-related clones infiltrated the inflamed tissue (159/159) (**Supplementary Fig.11c**), most 96.2% (153/159) were also present in normal tissue (**Supplementary Fig.11d**), indicating TLS-primed clones can migrate without inducing inflammation. Instead, disease activity correlated with enlarged plasma cell aggregates in the inflamed tissue (**Figure 4a, Supplementary Fig.10d)**, which contained the highest proportions of disease-related clones across IgH, IgK, and IgL families (**Figure 4e**). These findings underscore the critical role of plasma cell aggregates in driving inflammation in IBD.

To identify the origins of disease-related clones, we clustered 117 IgL clones from inflamed tissue into 18 families based on CDR3 sequence similarities (**Figure 4f-g**), classifying 70 as mucosal clones and 47 as disease-related clones. Of these, 61 were germline clones, and 56 were mutated clones (**Figure 4h**). Mucosal clones had a lower proportion of mutated clones (24.3%) compared to disease-related clones (83.0%), (**Figure 4h**), suggesting enhanced somatic hypermutation drives disease-related clone emergence. Lineage tracing using minimal Levenshtein distances **(Figure 4i)** revealed that 79.5% of mutated clones originated from top-expanded mucosal germline clones, while 17.9% could not be traced (**Figure 4j-k, Supplementary Fig.11e)**. These findings indicate that abnormal somatic hypermutation of clonal mucosal B/plasma cells in the lamina propria may contribute to IBD, aligning with evidence of microbiome-targeting antibodies in inflamed colonic plasma cells ^49^. Collectively, these data highlight the application of Stereo-XCR-seq for tracing lymphocyte lineages of autoimmune disease.

## Discussions

While single-cell analyses of TCR and BCR clonality and transcriptomics have advanced our understanding of inflammatory response, methods to apply this information *in situ* remain limited. Here, we developed Stereo-XCR-seq, enabling immune repertoire profiling at 500 nm resolution with key innovations. Stereo-XCR-seq achieves single-cell resolution and assigns CDR3 sequences to segmented cells, providing a precise definition of clonal expansion and enabling the exploration of lymphocyte clonal activities in micro-lymphoid aggregates (radius<100μm). To ensure unbiased T/BCR enrichment, we introduced sscirPCR, which targets the constant region instead of the variable region, overcoming primer specificity issues and biases seen in prior methods. This strategy efficiently retrieves T/BCR reads from Stereo-seq cDNA libraries and can be extended to other RNA-seq platforms with known adaptor sequences, facilitating immune repertoire analysis from retrospective cDNA libraries. Additionally, Stereo-XCR-seq uniquely provides paired TCR and BCR chains within the spatial transcriptome, though its pairing rate remains lower than 5’ scRNA-seq ^50^. Furthermore, by incorporating a short-read sequencing strategy, Stereo-XCR-seq improves cost-efficiency and scalability, making high-throughput immune repertoire analysis more accessible.

In the human specimens analyzed in this study, Stereo-XCR-seq enabled the identification of two distinct types of lymphoid aggregates - TLSs and plasma cell aggregates. These aggregates exhibited unique characteristics, including differences in the expression of TLS markers (*CXCL13*, *CR2*, *MS4A1,* and *CCL19*), cellular composition, and B/plasma cell clonal activities. While TLSs are known for priming antigen-specific T and B cells, plasma cell aggregates displayed strong CSR activity in BCRs. In mucosal tissues from IBD patients, we observed abnormally high proportions of somatically mutated clones compared to normal tissues, suggesting that plasma cell aggregates may also contribute to antigen-binding affinity maturation. Moreover, our findings suggest that somatic hypermutation and CSR can occur independently in both TLSs and plasma cell aggregates. Although our current dataset is limited in sample size and may not fully reflect the diversity of lymphoid aggregates in tumor or mucosal tissues, this study highlights the complexity of defining and stratifying lymphoid aggregates based on cellular traits, clonal activities, and transcriptomic profiles. Moving forward, large-scale pan-cancer or pan-disease studies will be essential to expand these insights. Nonetheless, our work demonstrates the potential of Stereo-XCR-seq as a transformative tool for immune-related studies, providing a framework for categorizing lymphoid aggregates across tissues and diseases.

A limitation of this methodology is the low pairing rate, which can make it challenging to define clones using paired chains in some tissues. To address this, we defined clones using single CDR3 chains instead, ensuring that not-expanded clones with few T/BCR reads were not omitted. In general, the pairing rate for TCRs was lower than for BCRs, likely due to the lower mRNA abundance in T cells. Increasing sequencing throughput could help improve the pairing rate. Additionally, like other TCR or BCR analysis methods, Stereo-XCR-seq cannot determine the antigen-specificities of clones. Incorporating methods such as fluorochrome-conjugated peptide-MHC (pMHC) staining could help locate antigen-specific T cells and identify their TCRs, as the juxtaposition of antigen, MHC, and TCRs is essential for T cell activation and clonal expansion ^51^. Given extensive clinical studies demonstrating the potential of improving adoptive T cell transfer therapy through antigen-targeting capabilities ^8, 52, 53^, we aim to further develop our platform to advance studies on TCR-antigen specificity in the future.

Sequencing-based and image-based spatial transcriptomics are two distinct parallel technologies to deliver similar and complementary measurements of gene expression *in situ* ^54^. Sequencing-based technologies keep the intact mRNA through polyA hybridization and *in situ* reverse transcription, which is essential to read VDJ sequences. Because of this, recently published tools that retrieved XCR reads to study spatial immune repertoire, including this study, are all based on sequencing-based technologies^12, 14, 15, 55^. Image-based spatial transcriptomics at subcellular resolution rely on complementary hybridization and signal enhancement by either *in situ* multiplexed hybridization or rolling circle replication ^56–63^. In comparison to other image-based methodologies, *in situ* sequencing exhibits potential application in diverse VDJ sequencing due to its untargeted nature, while this potential is restricted by the short read length ranging from 5 to 30 bases ^54, 57^. Recent improvement by incorporating *ex situ* sequencing extends the read length to 76 bases in Expansion Sequencing ^58^, still unmet for reading XCR. In the future, further elongation of the read length might substantiate the application of *in situ* sequencing in spatial immune repertoire investigations.

We envision that Stereo-XCR-seq could have broad applications in translational studies of immune-related diseases, particularly those involving abnormal plasma cell expansion. Disease-associated B/plasma cell expansions have been observed in atopic dermatitis ^64^, rheumatoid arthritis ^65^, and lupus nephritis ^66^, highlighting a strong correlation between antibody secretion activity and disease progression. While identifying autologous antigens remains challenging, Stereo-XCR-seq enables the identification of disease-associated T/BCR sequences, creating opportunities for paratope-targeting drug design. By narrowing down potential targets through the detection of aberrantly expressed antibodies, Stereo-XCR-seq can aid structure-based virtual screening in drug discovery. Furthermore, simulating antigen-binding affinity maturation - a natural structure-mimicking process in B cells – might inspire advancements in docking scoring methods, offering a promising path for therapeutic innovation.

## Methods

### Tissue access and process

#### Colorectal cancer cell lines and orthotopic murine model construction

The MC38-OVA orthotopic model was provided by Cyagen Biosciences with approval by the ethics committee of BGI Research (BGI-IRB A23027). MC38-OVA cells were cultured in Dulbecco’s modified Eagle’s medium (DMEM; Gibco, New York, USA) supplemented with 10% fetal bovine serum (FBS, HyClone, Massachusetts, USA) and 1% penicillin/streptomycin (Gibco, USA) in a 37°C humidified chamber with 5% CO2. A commercial mycoplasma detection kit (Vazyme, Cat# D101-01) was used for monthly contamination testing of the cell lines. A commercial mycoplasma detection kit (Vazyme, Cat# D101-01) was used for monthly contamination testing of the cell line. 5*10^6 cells were implanted into the cecum of the C57BL/6J mice. The mice were sacrificed by carbon dioxide suffocation 14 days after tumor cell implantation and the solid tumor tissues were collected. The tumors were rinsed twice using precooled PBS and wiped using Kimwipe tissue, then immediately embedded in Tissue-Tec OCT (Sakura, 4583) on dry ice and stored at -80°C.

#### Human samples collection

Two human cancer specimens were kindly provided by Peking University People’s Hospital. Two human mucosal biopsies samples were kindly provided by Zhejiang University School of Medicine. Two inflammatory bowel disease samples were collected from the normal and diseased tissue sites of the same patient. All samples were collected from untreated donors after either curative surgical resection or enteroscopic examination. The tissues were rinsed twice using precooled PBS and wiped using Kimwipe tissue, then immediately embedded in Tissue-Tec OCT (Sakura, 4583) on dry ice and stored at -80°C. This study was done in accordance with the Declaration of Helsinki. The protocol was reviewed and approved by The Institutional Review Board of BGI Research (BGI-IRB24066, BGI-IRB24012), Peking University (IRB00001052-24061) and Zhejiang University School of Medicine (IR2023396). The clinical information and specimens were collected with written informed consent from all donors.

### Preparation and sequencing of Stereo-seq library

The fresh-frozen tumor tissues were then transferred on dry ice to BGI Research for RNA Integrity examination. Tissue section was performed using Dakewe Cryostat Microtome (6250) at -20℃. 10∼15 50μm slices were collected for RNA extraction and integrity examination and a following 5μm slice was sectioned for H&E staining to determine the morphology of the specimens. Tissues with RIN value over 6.5 and desired FOVs were proceeded to Stereo-seq. After quality control, the specimens were serially sectioned at the thickness of 5μm (slice 1,2)-10μm (slice 3)-5μm (slice 4,5). Slices 2 and 4 were stained using H&E staining kit (Beyotime, C0105S) and scanned using Motic EasyScan System at 10x objective lens. Slices 1 and 5 were preserved in - 80℃ for further use. Slice 3 was flattened and attached to Stereo-seq chip, dried at 37℃ for 4 minutes, and then fixed in absolute methanol at -20℃ for 30 minutes. After fixation, the chip was processed to nucleus staining using Qubit™ ssDNA Assay Kit (ThermoFisher, Q10212). The chip was scanned using Motic EasyScan System at 10x lens to detect signal of ssDNA and washed with 0.1x SSC solution before permeabilization. Tissue was then permeabilized for mRNA precipitation and probe capture at 37℃, PH 2.0 for 5 minutes. *In situ* reverse transcription (42℃,1.5 hours) and tissue removal (55℃,1 hour) was performed in incubator. After that, CID and UMI co- barcoded cDNA was released, collected, and amplified by PCR for Stereo-seq library construction. The amplified product was proceeded to Stereo-XCR-seq enrichment. The library of Stereo-seq was sequenced using MGI DNBSEQ-T10 according to the manufacturer’s protocol.

### T and B cell receptors enrichment strategy from Stereo-seq library

To retrive the TCR and BCR transcripts from the amplified Stereo-seq library, we added ligating elements to Stereo-seq library by nested PCR using KAPA HiFi Hotstart Ready Mix (Roche, KK2602). 20ng purified dsDNA from the first round was input as the template and amplified using primer LE-F and primer LE-R (**Supplementary Table1**). The program of nested PCR was set as:

1. Denaturation: 95℃ for 5 minutes for denaturation. Heat lid was set to 105 ℃.
2. Amplification: 95℃ for 20 seconds, 50℃ for 20 seconds, 72℃ for 3 minutes. Repeating for 15 cycles.
3. Extension: 72℃ for 5 minutes.

All PCR products were purified using the 0.6x DNA clean Beads (VAHTSTM, N411-03) and quantified by Qubit^TM^ dsDNA Assay Kit (Thermo, Q32854).

400ng modified cDNA library was mixed with 20μM XCR splint oligo (**Supplementary Table1**) to generate the sscirDNA as followed:

1. Denaturation: 95℃ for 5 minutes. Heat lid was set to 105 ℃.
2. Splint-oligo mediated sscirDNA transformation: rapid cooling at -20 ℃ for 10 minutes.
3. Ligation: adding T4 DNA ligase and ligating buffer (BGI, Cat No. LS-EZ-E-00008O) according to the instructions. Incubating at 37 ℃ for 1 hour. Heat lid was set to 37 ℃.
4. ssDNA and double-stranded DNA (dsDNA) degradation: adding exonuclease I and III (BGI, Cat No.LS-EZ-E-00010P, Cat No.LS-EZ-E-00011P) and incubate at 37 ℃ for 30 minutes. Heat lid was set to 37 ℃.

Next, the sscirDNA was purified using 1.5X PEG32 beads (BGI, Cat No. L054) and input as the template and amplified using T/BCR constant primers (**Supplementary Table1**) for XCR enrichment. The enrichment was achieved by 2 rounds PCR. For the first round, 40ng sscirDNA was input as the template and amplified using constant primer 1-F and constant primer 1-R at a concentration of 0.5 μM. For the second round, 20ng purified dsDNA from the first round was input as the template and amplified using constant primer 2-F and constant primer 2-R at a concentration of 0.5 μM. Each T/BCR chain was processed respectively. The programs of 2 rounds of PCR were both set to:

1. Denaturation: 95℃ for 5 minutes for denaturation. Heat lid was set to 105 ℃.
2. Amplification: 98℃ for 20 seconds, 50℃ for 20 seconds, 72℃ for 3 minutes. Repeating for 15 cycles.
3. Extension: 72℃ for 5 minutes.

All PCR products were purified using the 0.6x DNA clean Beads (VAHTSTM, N411-03) and quantified by Qubit^TM^ dsDNA Assay Kit (Thermo, Q32854).

### Long-read library construction and sequencing

The purified enrichment products by constant primer 2 PCR were used as input into CycloneSEQ library preparation. Each T/BCR chain was proceeded respectively. The starting mass of long-read library construction was 1 μg. At least 2 library was constructed for each chain of each specimen. We used CycloneSEQ library kit(H940-000001-00) to carry out end-repaired and sequencing adapter ligation by following the commercialized instructions. The resulting products were purified using 0.6x DNA clean Beads (VAHTSTM, N411-03) and eluted in 30μl elution buffer. 1 μl was extracted for concentration measurements by Qubit^TM^ dsDNA Assay Kit (Thermo, Q32854). The libraries were then sequenced using the CycloneSEQ platform^19^.

### Short-read library construction and sequencing

Sequencing elements were then added to the enriched XCR libraries by nested PCR. 50ng XCR libraries of each chain was input as the template and amplified using sequencing adapter primers (Read1 and Read2, **Supplementary Table1**) at a concentration of 0.8 μM. The program of nested PCR was set to:

1. Denaturation: 95℃ for 5 minutes for denaturation. Heat lid was set to 105 ℃.
2. Amplification: 98℃ for 20 seconds, 65℃ for 20 seconds, 72℃ for 3 minutes. Repeating for 15 cycles.
3. Extension: 72℃ for 5 minutes.

All PCR products were purified using the 0.7x DNA clean Beads (VAHTSTM, N411-03) and quantified by Qubit^TM^ dsDNA Assay Kit (Thermo, Q32854).

For each specimen, we generated an XCR mix for each specimen using equal mass of the enrichment products of each chain. 60ng XCR mix was used to generate DNB using DNBSEQ one step make DNB kit (MGI, 1000020563) supplemented with 20μM PE150-splint oligo (**Supplementary Table1**). The DNB was loaded to sequencing chip of PE150 sequencing kit (MGI, 1000012555) and sequenced using MGISEQ-2000 instrument. The sequencing primers in the kits were replaced by XCR sequencing primer, including XCR sequencing primer 1, 2 and XCR MDA primer (**Supplementary Table1**). Sequencing program of Stereo-XCR-seq was set to: Read1(1-150bp, dark reaction 1-20bp), Read2(151-302bp, dark reaction 152-181bp).

### Stereo-seq raw data analysis

Fastq files were generated using a MGI DNBSEQ-T10 sequencer. Stereo-seq CID and MID are contained in the read 1 (CID: 1-25 bp, MID: 26-35 bp) while the read 2 consist of the cDNA sequences. The complete processing of Stereo-seq raw data was previously described ^23^ and now packed up as integrative pipelines on DCScloud platform (https://cloud.stomics.tech/#/login). We used spatial_RNA_visualization_v5 to ran Stereo-seq data for all samples for workflow consistency. The gem file, tissue mask, barcode whitelist (h5 file) was downloaded through the platform.

### Long-read XCR data processing

Split: We used the edlib Python package (https://github.com/Martinsos/edlib) to identify split element 1 “ATGGCGACCTTATCAG” (LD ⦤ 3), split element 2 “GCCATGTCGTTCTGTGAGCCAAGGAGTT” (minimum LD ⦤ 5), and element 3 “TTGTCTTCCTAAGAC” (the fixed sequence, minimum LD ⦤ 3) in each long read. We then checked the location of each split elements on each read. Only the LRs containing the combination of all split elements in the order as 1-2-3 were retained. According to the library structure, each raw read was split into a new read1 (including CID barcode and UMI) and a new read 2 (insert sequence) fq file with corresponding read ID. Only reads with CID barcode lengths between 20-30bp and insert sequences ⦤ 10,000bp were kept for downstream analysis.

Alignment and assembling: The new read 2 fq files were fed to MIXCR (version 4.6.0)^67^ for VDJ alignment with the parameter set to --preset generic-ont. The aligned reads were used for CDR3 assembling using the command - OassemblingFeatures=‘CDR3’ -OseparateByJ=true -OseparateByC=true - OseparateByV=true -OminimalQuality=5. The assembled CDR3 clone reads were exported using the exportClones and exportAlignments commands. Only clone reads were kept for the coordinate mapping.

Coordinate mapping: The tissue masks were generated using workflow spatial_RNA_visualization_v5 on DCS cloud (https://cloud.stomics.tech/#/dashboard). All the coordinates inside the tissue mask were defined as in-tissue coordinates. For each sample, we extracted the correspondence barcodes of all in-tissue coordinates from the Stereo-seq whitelist h5 files. We then performed k-mer splitting (k=5) and construct minhash vectors using the datasketch (https://github.com/ekzhu/datasketch) for each whitelist barcode. Thus, we built a locality-sensitive hashing (LSH) forest index for all whitelist barcodes. In the same way, we constructed minhash vectors in the same manner for the CID of each sample (the sequenced barcode). Then we performed approximate nearest neighbor (ANN) search to obtain the 10,000 whitelist barcodes closest to each sequenced barcode. Further, we used the edlib to screen the unique matches between whitelist and sequenced barcodes. Only matches with LDs <=4 were kept.

Filtering: Reads with CDR3 lengths between 5 and 30 amino acids were retained. The coordinates supported by only 1 read were discarded as low-fidelity coordinates. Subsequently, we calculated the inter-UMIs LDs of the same coordinate and grouped UMIs with a LDs of <=2 into the same UMI cluster. Only reads supported by the dominant UMI clusters were retained for XCR-metadata construction.

### Short-read XCR data analysi**s**

Split: We used the edlib Python package to search element 3 “TTGTCTTCCTAAGAC” (the fixed sequence, minimum LD ⦤ 1) in each raw short read 2 fq file. The 25bp sequences following the fixed sequence and the 10bp sequences preceding it were picked up as the CID barcode and the UMI respectively. The CID and UMI were combined and renamed as a new read 1 fq file (containing the CID and UMI). The raw short read 1 fq files were renamed as a new read 2 fq file (containing the insert sequences).

Coordinate mapping: The new read 1 file were aligned to the Stereo-seq whitelist h5 files using ST_barcodemap (https://github.com/STOmics/ST_BarcodeMap). Parameters were set to --mismatch 1 --umiStart 25. The reads failed in barcode mapping were discarded.

Alignment and assembling: The mapped new read 2 fq files were fed to MIXCR (version 4.6.0) ^67^ for VDJ alignment with the parameters set to -p rna-seq - OallowPartialAlignments=true -OsaveOriginalReads=true - OvParameters.geneFeatureToAlign="VTranscriptWithout5UTRWithP". The aligned reads were used for CDR3 assembling using the command - OassemblingFeatures=’CDR3’ -OseparateByJ=true -OseparateByV=true. The assembled CDR3 clone reads were exported using the exportClones and exportAlignments commands.

### XCR-metadata construction

For both short-reads(SRs) and long-reads(LRs), only reads with both coordinates and CDR3s were retained. XCR-metadata was constructed with the following information extracted from VDJ alignments, assembling and coordinate mapping: ReadID, alignVHits, alignDHits, alignJHits, alignCHits, cloneID, CDR3aa, CDR3nt, isotype, priisotype, func (functional clone or not), CID, UMI, x coordinate, y coordinate, location, readtype (SR or LR), CDR3@isotype, amendedCHits (SR only) and amendedCDR3@isotype (SR only). Among the above information, alignVHits, alignDHits, alignJHits, alignCHits, cloneID, cdr3aa, cdr3nt, isotype, priisotype and func were directly extracted from the VDJCA file. CID, UMI, x coordinate, y coordinate and location was extracted from the barcode mapping results. Readtype (SR or LR) and CDR3@isotype (combining the CDR3 and isotype information) was manually added. For those CDR3s detected by SR, C region could be missing due to the restriction of read length, while the UMI and CID information was retained. To address this, we screened the LRs using the UMI and CID. The C region and isoform information screened in the LRs were added to SR reads with identical UMI and CID. Of note, the CDR3s only supported by LRs were discarded.

### Cell segmentation and XCR pairing

The cell segmentation pipeline was previously described in detail ^20^. In the first step, we used the gene expression matrix at a resolution of bin1 (500nm) to generate the mRNA.png. The UMI counts in each bin1 spot were summed and indicated by grey value. The UMI counts were then written as an mRNA.png file using opencv2 (https://github.com/opencv/). The mRNA.png was opened up using photoshop and set as the background. All track lines of the mRNA.png were highlighted manually. The nucleus staining image was imported as the above layer and an editable object. We then adjusted the contrast and brightness of the nucleus staining image so that every track line in this layer could be clearly seen as the mRNA.png. The length-width ratio of the nucleus staining image was both fine-tuned to correct the lens distortion. For registration, we manually adjusted the angle and zoom-in ratio until all the track lines on the mRNA image and on the nucleus staining image were matched without visible shifts. Afterwards, all the adjustments of the contrast and brightness were restored. The resulting file was exported as the registered image.

The registered image was then imported to CellposeV2 ^21^ using "File->Load image (*.tif, *.png,*jpg) " function. We used default parameters in Views, Drawing, Segmentation and Image Saturation settings. We cropped the raw images to different sizes for segmentation training. Each cropped image contained 10-30 cells. All cropped images were imported, automatically segmented and manually corrected and training respectively. We used function "Models->Train new model with image+masks in folder" for training. The parameters of training mode were set as following: initial mode; chan to segment (0:gray); chane (0:none); learning rate: 0.1; weight decay (0.0001); n epochs (100). We trained 10-20 cropped image per sample. After training all cropped images, the trained model was employed for the segmentation of the registered image. The resulting file was saved as an *_seg.npy format and used to generate the cell masks. We then assigned each XCR reads to each cell according to the coordinates and the cells with both TCRα and TCRβ or both IgH and IgK/L were counted as paired cells.

### Hypermutation definition and family annotation

Only SR reads were used to evaluate hypermutation. We exported the V and J gene mutation from MIXCR pipeline using function “AllVHitsWithScore” and “AllJHitsWithScore”. We checked the V and J mutation of each CDR3 amino acid sequence. The CDR3 sequences with one or more mutated loci were defined as hypermutated CDR3s. Each CDR3 amino acid was defined as a clone.

To interpret the clonal family, we calculated the pairwise LDs of all the CDR3s of each isotype using Levenshtein package (https://rapidfuzz.github.io/Levenshtein). The inter- CDR3 LDs were transformed into a similarity matrix using function spatial.distance.squareform of Scipy package (https://scipy.org/). This similarity matrix was then fed to hierarchy.linage (method=‘ward’) to construct the hierarchical lineage tree. We then performed hierarchical clustering based on the minimum intra-class variance, by using the function cluster.hierarchy.fdluster with a threshold set to 20. The family tree plots were presented by Radial Tree package (https://github.com/koonimaru/radialtree).

### CSR definition and counting

For the NSCLC tumor, we picked up all the IgH CDR3 clones presented in the intratumoral TLS and checked whether different isoforms could be identified in the same CDR3 clones. The CSR events were defined as over 1 isoform could be detected in any CDR3s within each spatial cluster. The CSR events were counted respectively within each spatial cluster and presented as a squarified treemap using Squarify (https://github.com/laserson/squarify).

### lymphoid aggregates identification in ccRCC

We summed the normalized UMI counts of BCR genes (gene symbols starting with *IGL, IGK or IGH,* non-relavant genes were manually excluded such as *IGLON5*) in each bin50 SB. For denoising, we proceeded the SBs with top 85% BCR expression values as BCR-SBs using KDTree (https://scikit-learn.org/stable/). By setting k value to 10, we obtained the pairwise distances between each BCR-SB and its 10 neighboring BCR-SBs. To identify those gathering spots, we kept the BCR-SBs with small neighboring distances (top 20%, ranked in ascending order) and annotated them as lymphoid aggregates. These lymphoid aggregates were then processed using DBSCAN function (eps=100, min_samples=3) for clustering.

### Tumor border fitting curve construction

For the NSCLC tumor, we used KDTree function obtained the pairwise distances between each alveolar SB and its 5 neighboring alveolar SBs. The bins with distances ranked in the least 5% (ascending order) were discarded for denoising. Further, we used the function interp1d (Scipy package) to refill the alveolar region and performed convex hull computation using Concaveman (https://github.com/mapbox/concaveman). The convex hulls were smoothed by Gaussian smoothing using function gaussian_filter1d (sigma=10) and then annotated as the tumor border.

### Intra/peri-tumoral TLS definition

For the TLSs in the NSCLC tumor, we used KDTree function obtained the pairwise distances between each TLS SB and its 150 neighboring TLS SBs. The bins with distances over median distances were discarded for denoising. These TLS bins were then processed using DBSCAN function (eps=300, min_samples=5) for clustering. The TLSs located on the tumor border were annotated as peritumoral TLSs. The TLS located in the tumor microenvironment was annotated as intratumoral TLS otherwise.

### Deconvolution of NSCLC SBs

The scRNA-seq data of NSCLC was fetched from https://www.ncbi.nlm.nih.gov/geo/query/acc.cgi?acc=GSE148071 ^68^. The raw data was normalized and clustered using Scanpy (https://scanpy.readthedocs.io/en/stable/index.html). The cell clusters were manually annotated to 13 major clusters according to the expression genes of interests (Supplementary Figure 7c). Deconvolution of cell clusters in the spatial map was calculated using robust cell type decomposition (RCTD) ^69^ algorithm in R package spacexr-2.0.018 in combined with scRNAseq dataset. The raw counts of scRNA-seq (with annotation) and the Stereo-seq were input to construct the RCTD object. For Stereo-seq data, we set the minimal UMI count to 0 to avoid filing out any spots. Since the cell diameters typically ranged from 5 to 12 μm, the SBs may contain multiple cell types that might not be consistent across the spot. Therefore, we ran the RCTD in full mode, without restrictions on the number of cell types. In this study, the normalized probabilities of cell types were referred to as “RCTD frequencies”, which were exported and integrated into an h5ad objects for further visualization and analysis.

### Lineage tracing of IgH clones

For the IBD specimens, we firstly obtained the germline and mutated IgH CDR3s of the inflamed tissue. The germline clones could be also found in normal tissue were defined as mucosal ancestor clones, while the other germline clones were defined as immigrant ancestor clones. We then used KDTree to calculate the pairwise LDs between germline and mutated IgH CDR3s. The mutated CDR3s failed in lineage tracing (LD >=5 to all germline CDR3s) were termed as undefined clones. The mutated CDR3s successfully mapped to mucosal ancestor clones (LD<5 & the LD to t mucosal ancestor clones < the LD to immigrant ancestor clones) were defined as mucosal daughter clones. The mutated CDR3s successfully mapped to immigrant ancestor clones (LD<5 & the LD to immigrant ancestor clones < the LD to mucosal) were defined as immigrant daughter clones. The mutated CDR3s with equal LDs to immigrant and mucosal ancestor clones were defined as ambiguous clones.

### Shannon’s Index Calculation

We inferred the locoregional clonal diversity by calculating the Shannon’s index within each cluster of interests. We obtained the geometric center of each segmented cell and assigned them to spatial clusters. Then, we counted the absolute cell number of each CDR3 calculated the frequencies of each CDR3 as 𝑃_𝑖_ . The Shannon’s index was calculated using formula 1:

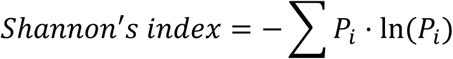

## Data and code availability

The raw sequencing FASTQ files of the spatial transcriptome and Stereo-XCR-seq data could be accessed on Genome Sequence Archive for Human (GSA-Human, BioProject accession number HRA009729). The processed matrix of Stereo-XCR-seq, gene expression matrix and barcode whitelist have been deposited into STOmicsDB ^70^ of China National GenBank Database ^71^ with accession number STT0000123. The external single cell RNA-seq data could be downloaded through the following links: https://www.ncbi.nlm.nih.gov/geo/query/acc.cgi?acc=GSE148071. Code used in this study has been uploaded to github (https://github.com/fengyu9481) for the reproducibility of this study. For any further inquiries for the use of the code or for the data accession, please send correspondence to.

## Acknowledgement

This work was supported by Shenzhen Key Laboratory of Single-Cell Omics (ZDSYS20190902093613831), National Natural Science Foundation of China (Grant No. 92374116), and Shenzhen Proof-of-Concept Center of Digital Cytopathology. This work was also supported by BGI Research, Shenzhen, BGI Research, Hangzhou and BGI Hangzhou CycloneSEQ Technology Co., Ltd. We thank China National GeneBank, BGI Research, Shenzhen for data storage, open use, and management.

## Author contributions

Methodology development: Yu Feng, Xiaojuan Zhan;

Data analysis and coding: Yi Liu, Young Li, Yixin Yan, Hui Zeng, Fan Zhu, Zhong Liu, Xinxin Li;

Sample collection and preparation: Zexian Zeng, Jinwen Yin;

Library construction: Yu Feng, Xiaojuan Zhan, Yanying Guo, Xiaoyu Chen, Rong Ma; Short reads sequencing: Xiaojuan Zhan, Yanying Guo, Xiaoyu Chen, Xuan Dong; Long reads sequencing: Yuliang Dong, Tao Zeng, Longqi Liu;

Pathological annotation: Jinwen Yin, Francis Ka-ming Chan;

Manuscript drafting: Yu Feng, Jingying Zhou, Wenwen Zhou, Haoran Tao, Zexian Zeng, Xun Xu, Chuanyu Liu, Yong Hou;

Review & editing: Zexian Zeng, Xubin Zheng, Francis Ka-ming Chan, Longqi Liu; Conceptualization and supervision: Young Li, Jingying Zhou, Zexian Zeng, Yu Feng.

