## Supplementary Figure for "Single cell resolved spatial immune repertoire unveils spatial heterogeneity of lymphoid aggregates in human immune disorders"


**Authors/affiliations:**

Xiaojuan Zhan[1,2,3*], Yi Liu[1,3*], Yanying Guo[2], Wenwen Zhou[1,2], Yixin Yan[3], Hui Zeng[1,2], Xuan Dong[3], Xiaoyu Chen[3], Rong Ma[3,4], Zhong Liu[3], Fan Zhu[1,3], Xubin Zheng[5,6], Xinxing Li[3], Jinwen Yin[7], Francis Ka-ming Chan[7,8,9,10], Chuanyu Liu[2,11,12,13], Longqi Liu[2,3,12] , Xun Xu[1,2], Yong Hou[1,2,14], Haoran Tao[2,15], Yuliang Dong[2,16], Tao Zeng[2,16], Young Li[3#], Jingying Zhou[2,15#], Zexian Zeng[17,18#], Yu Feng[2,3,12#]

[1]College of Life Sciences, University of Chinese Academy of Sciences, Beijing 100049, China

[2]BGI Research, Shenzhen 518083, China

[3]BGI Research, Hangzhou 310030, China

[4]College of Life Sciences, Northwest University, Xi'an 710069, China

[5]School of Computing and Information Technology, Great Bay University, Dongguan 523000, China

[6]Guangdong Provincial Key Laboratory of Mathematical and Neural Dynamical Systems, Dongguan 52300, China

[7]Zhejiang University Liangzhu Laboratory, 1369 West Wenyi Road, Hangzhou 311121, China

[8]Department of Cardiology of The Second Affiliated Hospital, School of Medicine, Zhejiang University, Hangzhou 310009, China

[9]State Key Laboratory of Transvascular Implantation Devices, Hangzhou 310009, China

[10]Heart Regeneration and repair Key Laboratory of Zhejiang Province, Hangzhou 310009, China

[11]Shenzhen Proof-of-Concept Center of Digital Cytopathology, BGI Research, Shenzhen 518083, China

[12]Shanxi Medical University - BGI Collaborative Center for Future Medicine, Shanxi Medical University, Taiyuan 030001, China

[13]Key Laboratory of Anti-Inflammatory and Immune Medicine, Ministry of Education, Anhui Medical University, Hefei 230032, China

[14]Shenzhen Key Laboratory of Single-Cell Omics, BGI Research, Shenzhen 518083, China

[15]School of Biomedical Sciences, The Chinese University of Hong Kong, Hong Kong SAR, 999077, China

[16]BGI Hangzhou CycloneSEQ Technology Co., Ltd, Hangzhou 310030, China

[17]Peking-Tsinghua Center for Life Sciences, Academy for Advanced Interdisciplinary Studies, Peking University, Beijing 100084, China

[18]Center for Quantitative Biology, Academy for Advanced Interdisciplinary Studies, Peking University, Beijing 100084, China

*These authors contributed equally to this work

**Correspondence:**

#### Supplementary fig 1

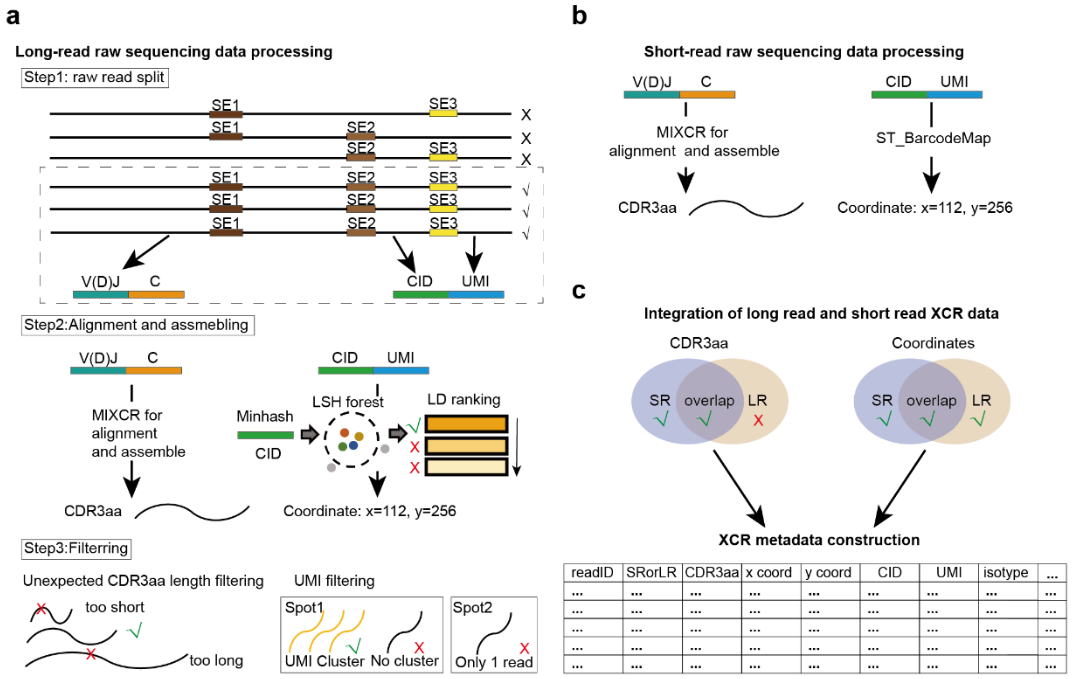


**Supplementary Figure 1.** ***The analytic pipelines of stereo-XCR-seq.***
**a.** Schematic diagram of long-read processing strategy. Raw read are split by identify split elements 1~3 (SE1~3) and retained reads that contain the combination of all split elements in the order of 1-2-3. Split raw reads into a new read1 (including CID barcode and UMI) and a new read 2 (insert sequence) fq file with the corresponding read ID (step1). The new read 2 is aligned and assembled using MIXCR, and clone reads are kept for coordinate mapping (step2). Filter reads with CDR3 lengths between 5 and 30 amino acids and coordinates supported by the dominant UMI clusters for XCR-metadata construction (step3). b. Schematic diagram of short-read processing strategy. Raw reads are split by identifying split element 3 in the short read 2 fq file to obtain coordinates (including CID barcode and UMI) for ST_BarcodeMap. The mapped reads are fed to MIXCR for alignment and assemble. **c.** The schematic diagram shows the construction of XCR metadata. Short-read (SR) and long-read (LR) reads with both coordinates and CDR3s are retained for XCR-metadata construction.

#### Supplementary fig 2

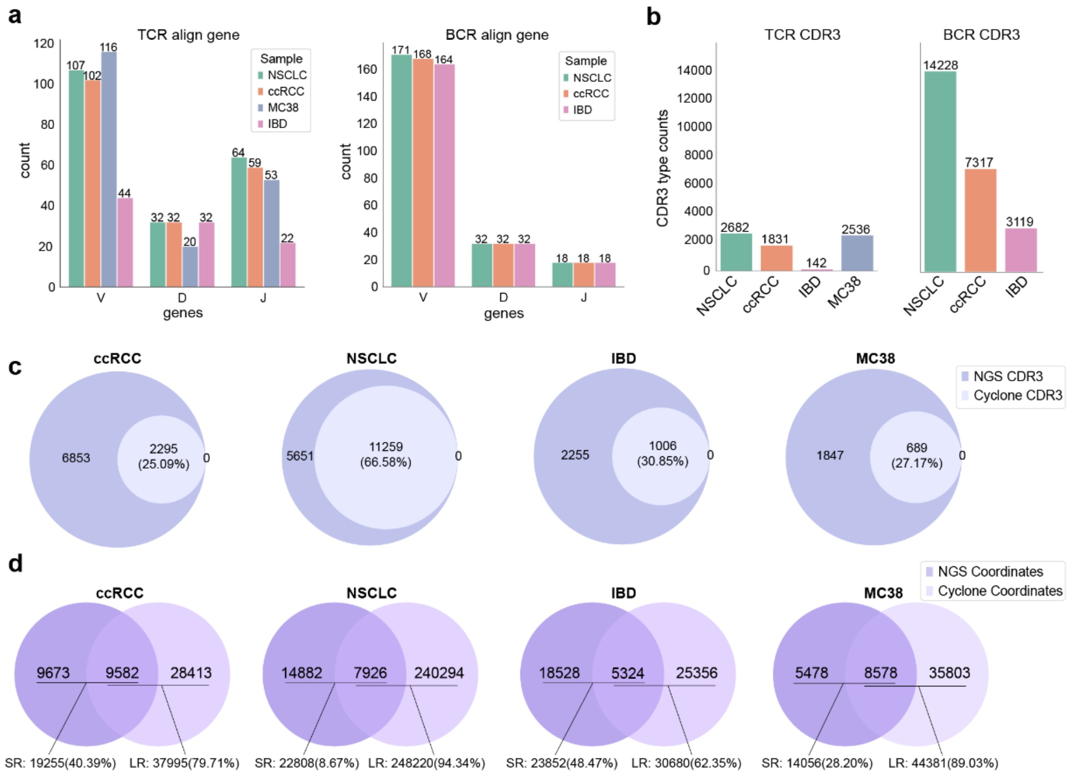


**Supplementary Figure 2.** ***Stereo-XCR-seq sample application and quality control.***
**a**. The bar charts show the number of V(D)J genes detected in TCR and BCR of each sample. The cell numbers are labeled above each bar. **b.** The bar plot shows the CDR3 clones of each sample. The numbers of clones are labeled above each bar. **c-d.** The Venn diagrams show the CDR3 types (**c**) and coordinates (**d**) detected by short-read (SR) and long-read (LR) sequencing, respectively.

#### Supplementary fig 3

a

| Technologies | Based on spatial transcriptome | Resolution | Enrichment approach | Capture primer/probe target | Bias in enrichment | Available enriched type | Available sequencing type | | Pair chains in cells |
|---|---|---|---|---|---|---|---|---|---|
| | | | | | | | Long reads | Short reads | |
| Stereo-XCR-seq | Stereo-seq | Single cell | sscirPCR | Constant region | Unbiased | TCR and BCR | TCR and BCR | TCR and BCR | TCR:α & β<br>BCR:IGH & L |
| Slide-TCR-seq [1] | Slide-seq | 10 μm | Mulitiplex PCR | Variable region | Biased | TCR | TCR | TCR | No |
| Slide-tags[2] | Slide-seq | 10 μm | Multiplex PCR | Variable region | Biased | TCR | No | TCR | postulative<br>TCR:α & β |
| Spatial-VDJ [3] | 10x Visium | 100 μm | Probe hybridization and Mulitiplex PCR | Constant region probes<br>Variable region TCR primers | Biased TCR<br>Unbaised BCR | TCR and BCR | TCR and BCR | TCR | No |
| SPTCR-seq [4] | 10x Visium | 100 μm | Probe hybridization | V(D)JC region | Unbiased | TCR | TCR | TCR | No |

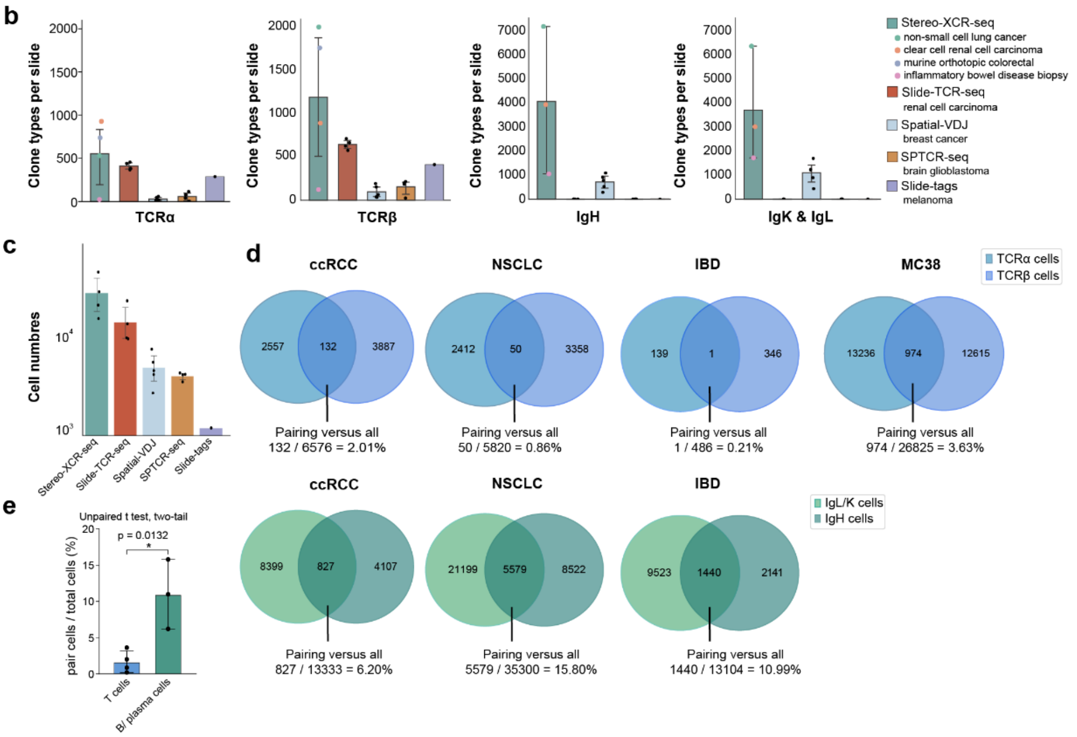


**Supplementary Figure 3.** ***Benchmarking of Stereo-XCR-seq for spatial immune repertoire profiling.***

**a**. The technology comparison of stereo-XCR-seq, Slide-TCR-seq [1], Slide-tags [2], Spatial-VDJ [3] and SPTCR-seq [4]. **b.** The bar chart shows the clone types detected in each slide using stereo-XCR-seq versus external datasets of non-lymphoid tissues. Each dot represents a slide. For stereo-XCR-seq, dots are colored by different sample types. Data are presented as mean±SD. N slides: stereo-XCR-seq =4, Slide-TCR-seq=4, Spatial-V(D)J=5, SPTCR-seq=4, Slide-tags=1. **c.** The bar chart shows the cell numbers detected in each technology. N slides: stereo-XCR-seq =4, Slide-TCR-seq=4, Spatial-V(D)J=5, SPTCR-seq=4, Slide-tags=1. Data are presented as mean±SD. **d.** The Venn diagrams show the TCRα/β and BCR H-L pairing cells. The overlapped area indicates the cells contained both TCRα/β or BCR H-L chains. Pairing rates are calculated and labeled below each Venn diagram. **e.** The bar plot shows the difference between the pairing rates of TCR and BCR. The comparison is calculated using two-tailed unpaired t-test. *, $p<0.05$. Data are presented as mean±SD.

#### Supplementary fig 4

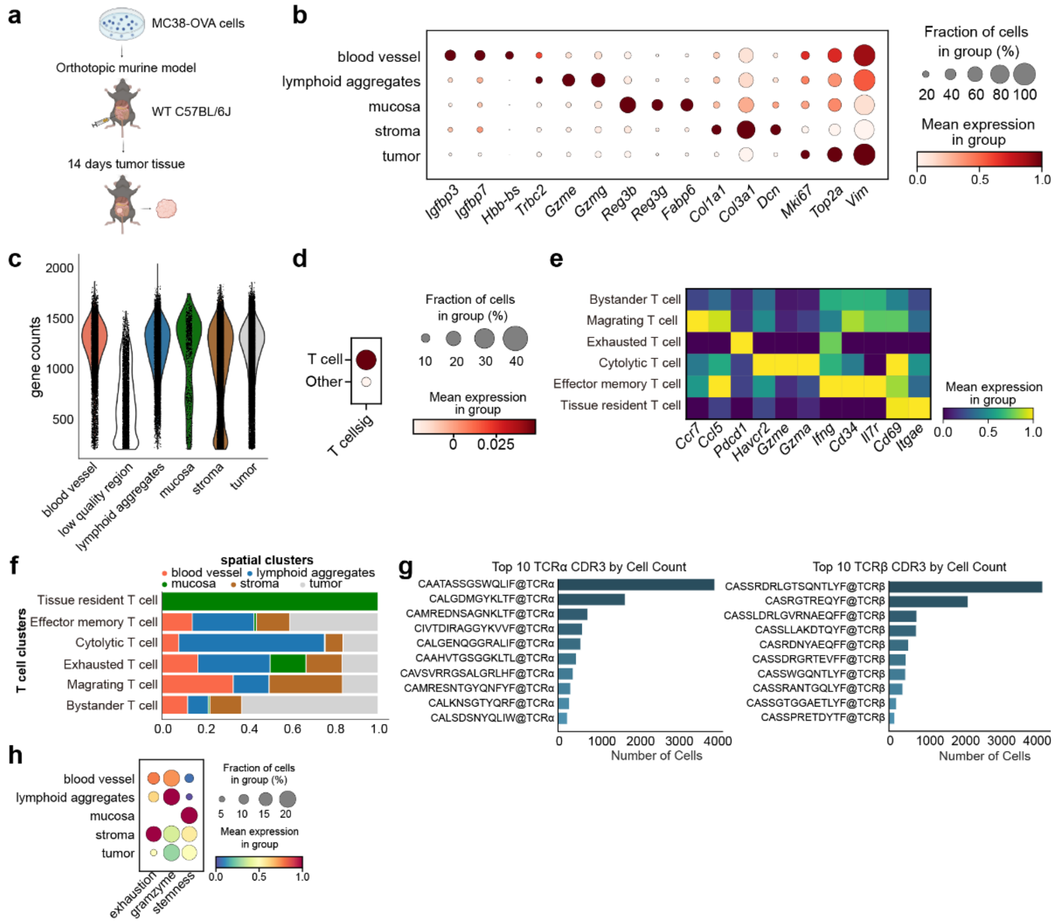


**Supplementary Figure 4.** ***The spatial transcriptome and spatial immune repertoire of MC38-OVA tumor.***
**a.** The schematic diagram shows the implantation and harvest of the orthotopic MC38-OVA tumor. **b**. The dot plot shows the expression of genes of interest in each spatial cluster (bin50). The dots are colored by mean expression and sized by the fraction of cells in each cluster. The expression of each gene is standardized by cluster for data presentation. **c**. The violin plot shows the gene counts per spot in each spatial cluster (bin50). Each dot represents a square bin. The violins are colored by spatial clusters. **d**. The dot plot shows the expression of T cell signature scores of segmented T cells (with either TCRα or β chain or both) and other segmented cells (without TCRα/β chains). The dots are colored by mean expression and sized by the fraction of cells in each cluster. **e**. The matrix plot shows the mean expression of the genes of interest of each T cell clusters. The matrixes are colored by mean expression. The expression of each gene is standardized by cluster for data presentation. **f**. Stacked bar plot shows the proportions of each T cell clusters in the indicated spatial clusters. The stacked bars are colored by spatial clusters. **g**. The bar plot shows the top 10 expanded TCRα and β clones. Bars are colored by ranking and lengthen by clonal sizes (number of cells). **h**. The dot plot shows the expression of genes of interest by the T cells located in different spatial clusters. The dots are colored by mean expression and sized by the fraction of cells in each cluster. The expression of each gene is standardized by cluster for data presentation.

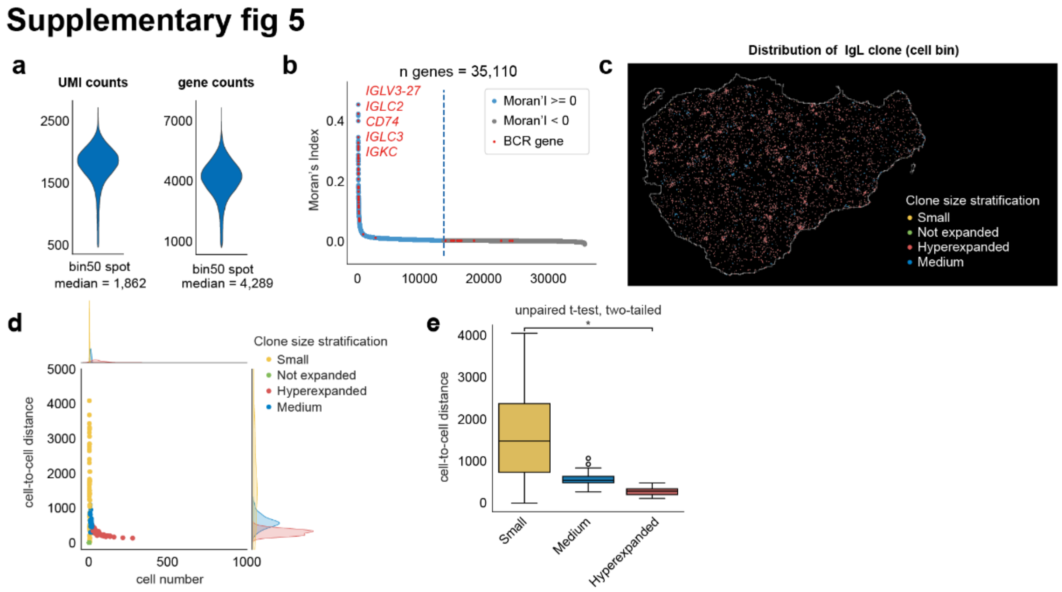


**Supplementary Figure 5.** ***The spatial distribution of clonal T cells and B/plasma cells in the ccRCC tumor***

**a**. The violin plots show the UMI counts (left) and gene counts (right) of the square bins (bin50) of the ccRCC tumor. The median values are labeled below each violin. **b**. The scatter dot plot shows the Moran's indices of the 35,110 genes detected by the spatial transcriptome. Each dot represents a gene, ranked by the Moran's indices value. The genes with a Moran's index above or equal to 0 are colored in blue and the BCR-related genes are highlighted in red. The dotted vertical line indicates the gene with a Moran's index equal to 0. **c**. The spatial plot shows the spatial distribution of the hyperexpanded (clone sizes$\geq$20), medium (6$\leq$clone sizes < 20) small (2$\leq$clone size < 6) and not expanded (clone size=1) IgL clones at a resolution of cell bin. The cells are colored by the clone size stratification. **d**. The scatter dot plot shows the nearest cell-to-cell distances of each IgL clone. Each dot represents a clone. The curve outside the graph indicates the distribution of cell number of each clone (above) and cell-to-cell distance (right side). Both scattered dots and curves are colored by clone size stratification. **e**. The box-whisker plot shows nearest cell-to-cell distances of each IgL clone crossing different clone stratifications. The boxes are colored by clone size stratification. The comparison is made by a two-tailed unpaired t-test. *, $p<0.05$. Data are presented as median ± IQR (25%~75%) with extreme values indicated by lower and upper bars. The outliers are indicated as hollow dots.

### Supplementary fig 6

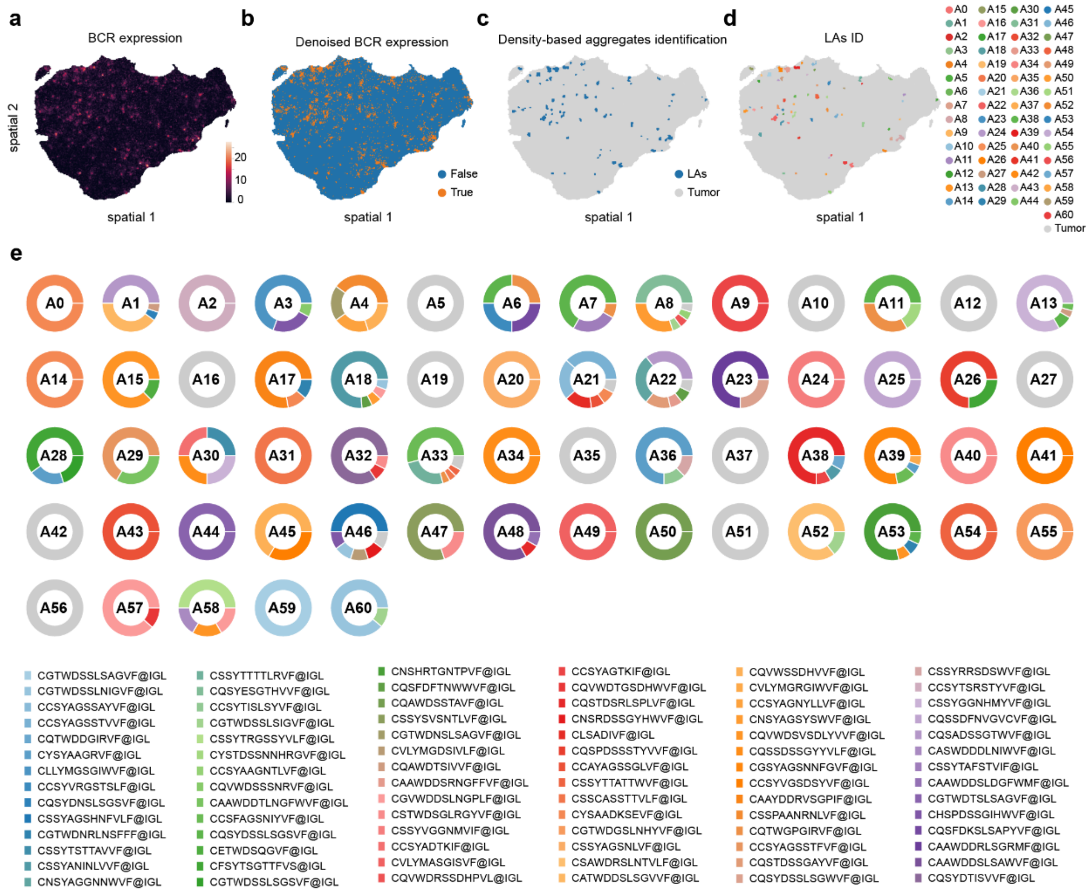


**Supplementary Figure 6.** ***The clonality of B/plasma cells in the ccRCC tumor.*** **a-d.** The spatial map shows the expression of BCR-related genes (**a**, bin50), denoised expression of BCR-related genes (**b**, bin50), and density-based LAs identification (**c-d**, bin50). In **d**, the LAs are colored by the numeric IDs. **e.** The donut plots show the cell proportion of each IgL clone across LAs. Each sector is colored by the CDR3 amino acid sequences, which are labelled below.

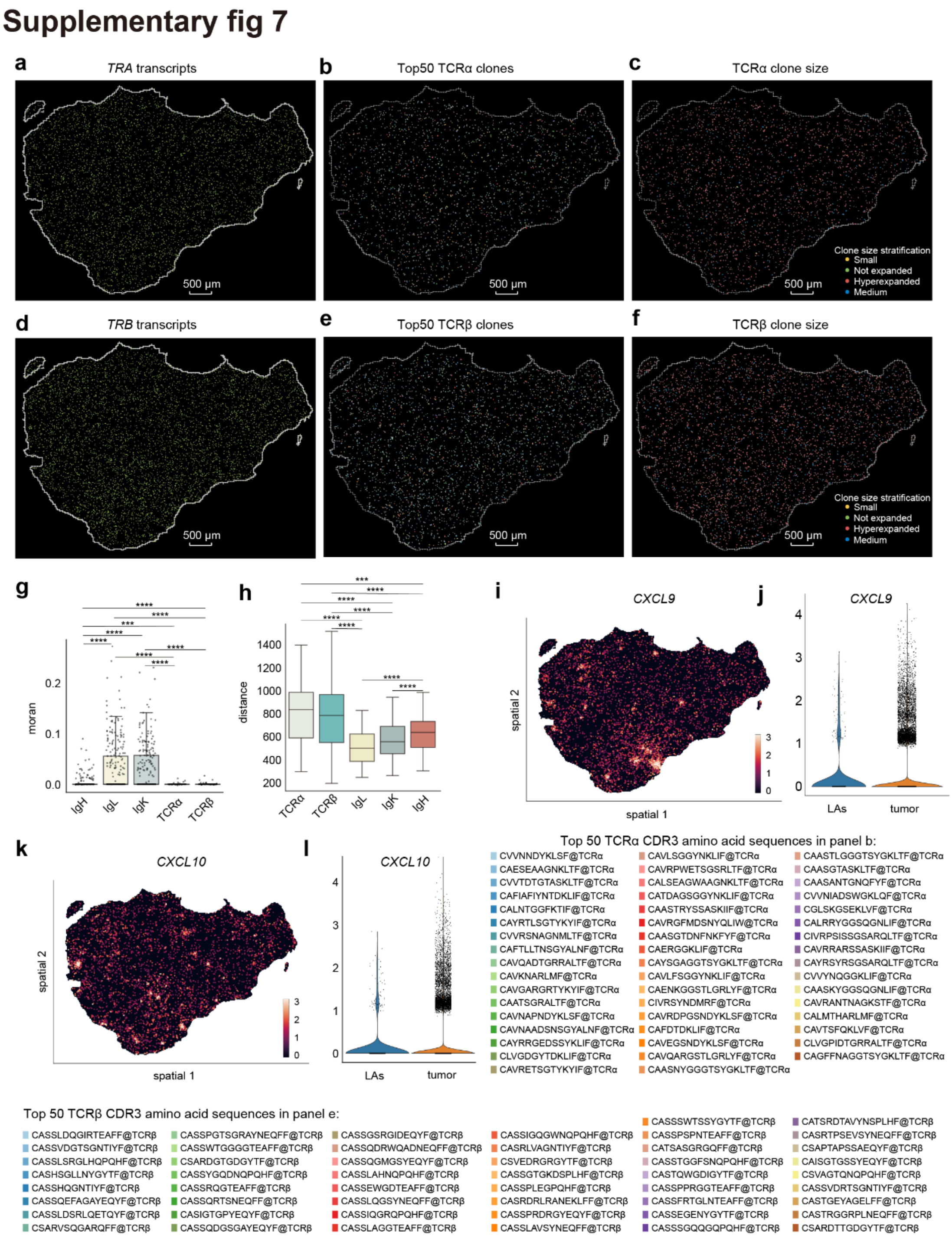


**Supplementary Figure 7.** ***The clonal expansion of T cells and B cells in the ccRCC tumor.*** **a-f.** The spatial plot shows the spatial distribution of *TRA* (**a**) and *TRB* (**d**) transcripts at a resolution of 500nm (bin1), the spatial distribution of the top 50 TCRα (**b**) and β (**e**) clones at a resolution of cell bin, the hyperexpanded (clone sizes≥20), medium (6≤clone sizes < 20) small (2≤clone size < 6) and not expanded (clone size=1) TCRα (**c**) and β (**f**) clones at a resolution of cell bin. Dots in **a &d** with at least one *TRA/TRB* transcript (**c**) or one TCRα or β clone read are colored in green. Cells are colored by CDR3 amino acid sequences in **b & e** and by clone size stratification. **g.** The box-whisker plot shows the Moran's indices of each type of receptor chain. Each dot represents a clone type. The comparison is made by a two-tailed unpaired t-test. *, $p<0.05$. Data are presented as median ± IQR (25%~75%) with extreme values indicated by lower and upper bars. **h.** The box-whisker plot shows the nearest cell-to-cell distances of each clone across different chains. The comparison is made by a two-tailed unpaired t-test. *, $p<0.05$. Data are presented as median ± IQR (25%~75%) with extreme values indicated by lower and upper bars. **i-l**. The spatial plots and violin plots show the expression of *CXCL9* (**i & j)** and *CXCL10* (**k & l**) in the ccRCC tumor. The spatial plots are presented at a resolution of bin50, with each dot colored by the normalized expression values. Each dot in the violin plot represents a bin50 spot.

#### Supplementary fig 8

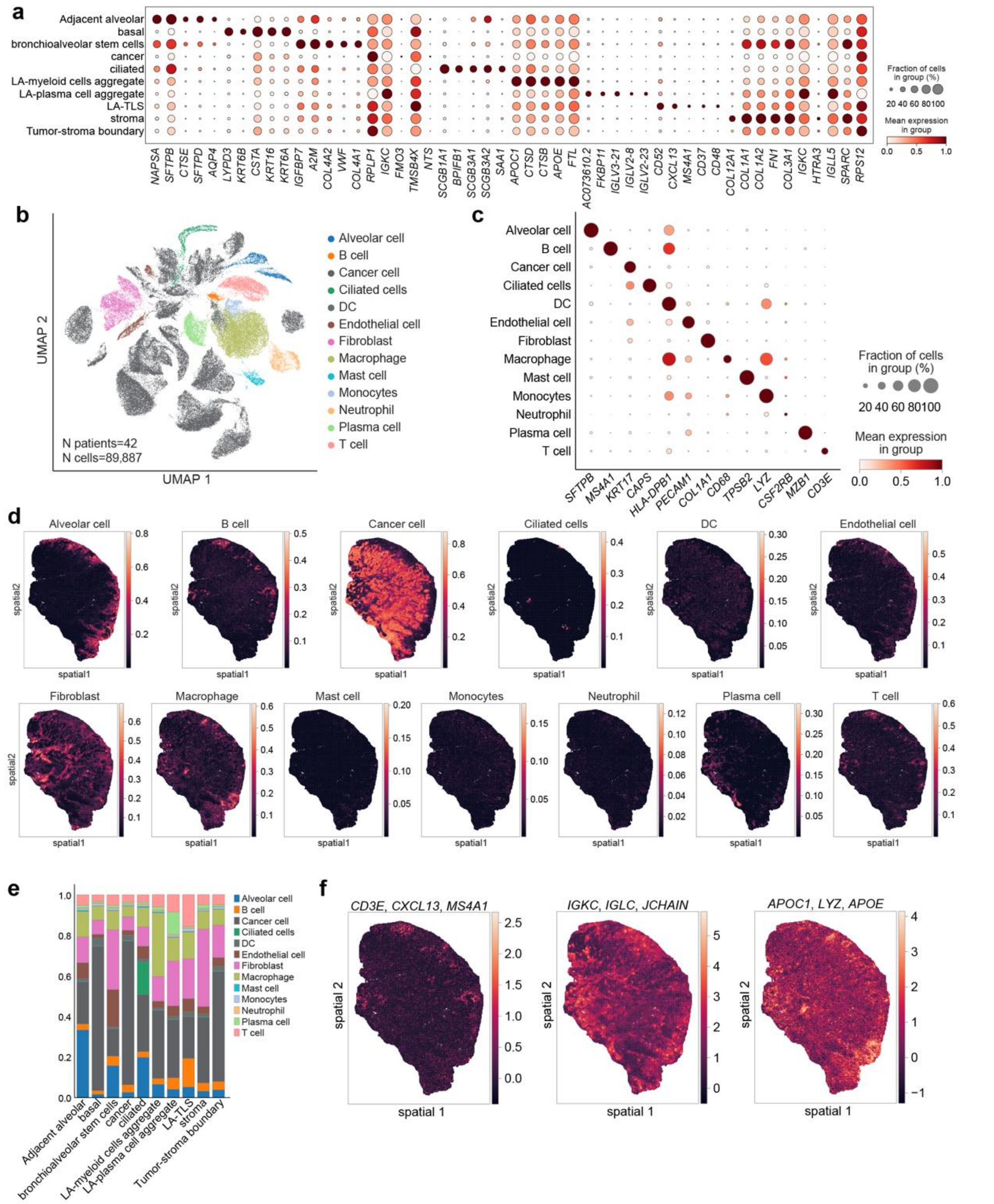


**Supplementary Figure 8.** ***The spatial transcriptome and annotation of the NSCLC.***
**a.** The dot plot shows the expression of DEGs in each spatial cluster (bin50). The dots are colored by mean expression and sized by the fraction of cells in each cluster. The expression of each gene is standardized by cluster for data presentation. **b.** The UMAP shows the external scRNA-seq dataset (GSE148071) [5]. Each dot represents a cell, colored by cell clusters. N patients=42, n cells=89,887. **c**. The dot plot shows the expression of intuitive markers in each single cell clusters. The dots are colored by mean expression and sized by the fraction of cells in each cluster. The expression of each gene is standardized by cluster for data presentation. **d**. The spatial plots show the distributive pattern of each cell type. The distributive probabilities are calculated by deconvoluting the spatial

transcriptome (bin50) using annotated scRNA-seq datasets (**b**). Each dot represents a square bin, colored by the distributive probability of the cells indicated above. **e**. The stacked plot shows the proportion of each cell type across each spatial cluster. The stacked bars are colored by single cell clusters. **f.** The spatial plots show the expression of TLS signature (left), plasma cell signature (middle), and myeloid cell signature (right) in the NSCLC tumor. Each dot represents a square bin (bin50), colored by expression level. The genes constituting the signatures are listed above each spatial plot.

#### Supplementary fig 9

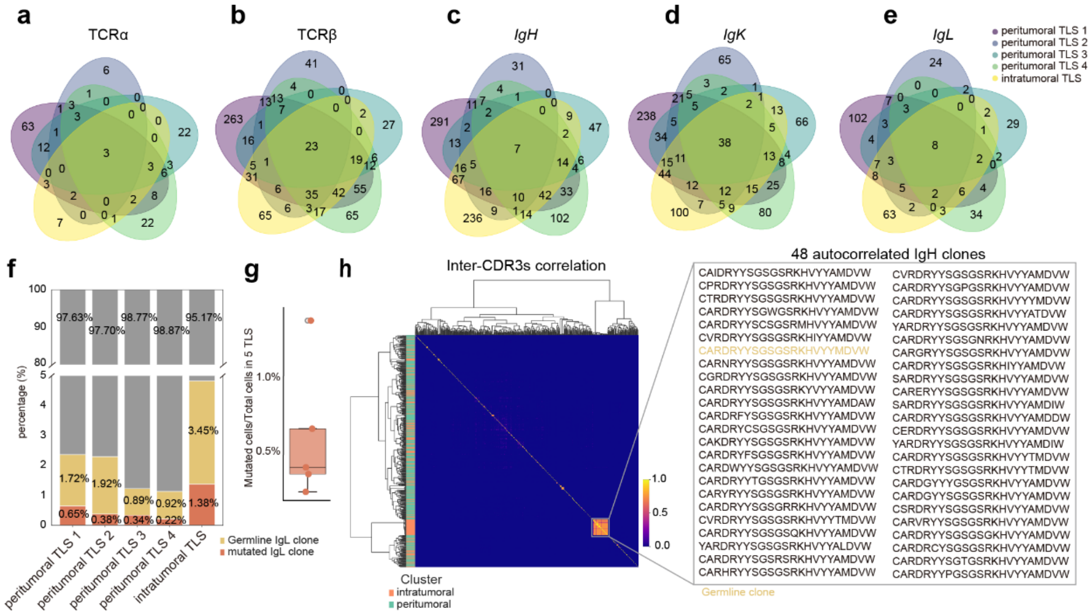


**Supplementary Figure 9.** ***The immune repertoire of the 5 geographically discrete TLSs.*** **a-e.** The Venn diagrams show the sharing of the clones of each chain. Each circle is colored by the TLS labeling as indicated on the right. **f.** The stacked plot shows the proportion of germline and mutated IgL clones in each TLS. **g**. The box-whisker plot shows the proportion of mutated B/plasma cells in the 5 TLSs. Data are presented as median ± IQR (25%~75%) with extreme values indicated by lower and upper bars. The outlier is indicated as a red dot. **h.** The heatmap shows the pairwise CDR3 similarities. Each matrix is colored by the inferred similarity. The colored lines on the left Y-axis are colored by the location of the TLSs. The tree plots on the left and the above are identical to indicate the lineage inference of the IgH clones detected in the 5 TLSs.

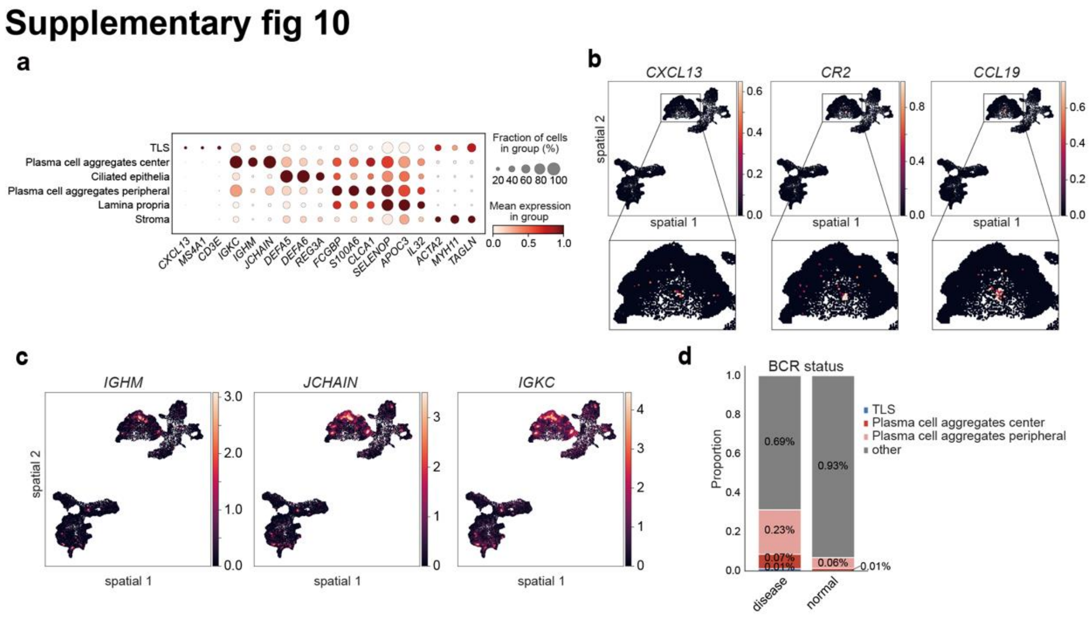


**Supplementary Figure 10.** ***The spatial transcriptome of the 2 biopsies from the IBD patient.***
**a.** The dot plot shows the expression of DEGs in each spatial cluster (bin50). The dots are colored by mean expression and sized by the fraction of cells in each cluster. The expression of each gene is standardized by cluster for data presentation. **b**. The spatial plot shows the gene expression of *CXCL13*, *CR2,* and *CCL19* in TLS of the inflamed IBD tissue. **c**. The spatial plot shows the gene expression of *IGHM*, *JCHAIN,* and *IGKC* in plasma cell aggregates of the inflamed IBD tissue. **d**. The stacked plot shows the area of lymphoid aggregates in inflamed tissue and normal tissue. The areas are counted using the number of square bins (bin50).

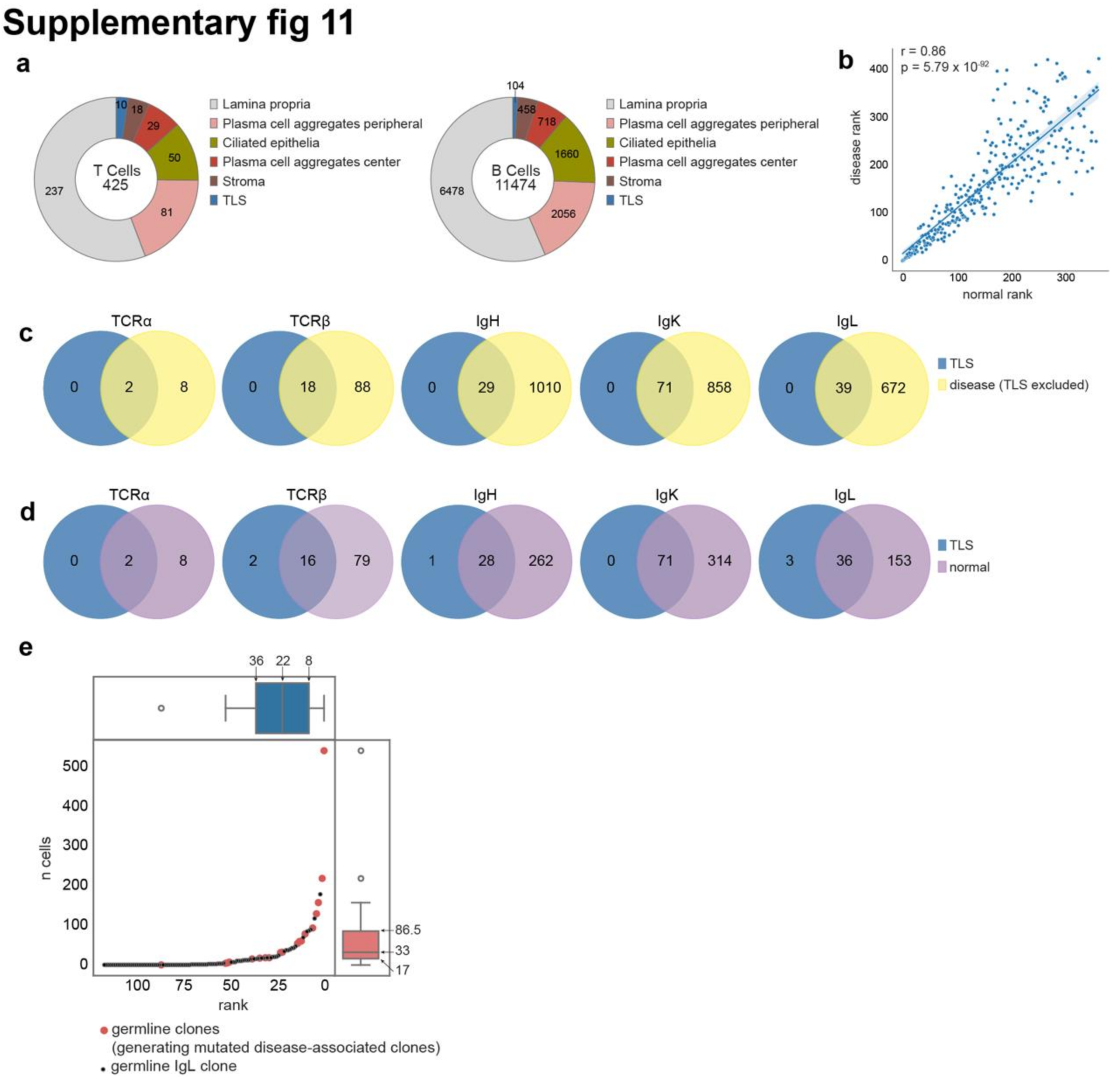


**Supplementary Figure 11.** ***The spatial immune repertoire of the 2 biopsies from the IBD patient.*** **a.** The donut plots show the distribution of T cells and B cells in the 2 biopsies from the IBD patient. **b.** The scatter plot and fitting line show the consistent clonalities of the shared clones in normal tissue (X axis) and inflamed disease tissue (Y axis). The P-value and r-value of the Pearson's correlation analysis are labeled above. **c-d.** The Venn plot shows the shared and unique CDR3 types of TLS and disease tissue (TLS excluded) (**c**) and of TLS and normal tissue **(d)**, respectively. **e.** The scatter dot plot shows the clonal size of IgL clones in the inflamed tissue. The germline clones that generated mutated disease-associated clones are highlighted as red enlarged dots. The distribution of the germline clones that generate mutated disease-associated clones is summarized as a box-whisker plot above (by ranking, blue) and on the right (by clone size, red) of the scatter dot plot. The box-whisker plot is presented as median ± IQR (25%~75%) with extreme values indicated by lower and upper bars. The outlier is indicated as a red dot.
